# Acute *LINE-1* overexpression rewires the transcriptome, proteome and secretome of primary human fibroblasts

**DOI:** 10.64898/2026.09.03.749265

**Authors:** Juan I. Bravo, Eyael Tewelde, Christina D. King, Nayana S. Tellakula, Birgit Schilling, Bérénice A. Benayoun

## Abstract

A large portion of eukaryotic genomes is composed of transposable elements, which are usually kept repressed in young, healthy cells but can be derepressed in response to cell senescence and organismal aging. *LINE-1 (L1)* is the most abundant TE in the human genome by percent coverage, and its derepression has been implicated in inflammaging responses. However, whether *L1* transcription itself is sufficient to drive aging-associated molecular phenotypes in healthy cells has not been extensively explored. Here, we leverage a multi-omic approach combining transcriptomics, proteomics, and secretomics of primary human IMR-90 fibroblasts transiently overexpressing a human *L1Hs* element in order to define the systems-level consequences of *L1* transcription. Intriguingly, transient *L1Hs* overexpression induced widespread remodeling of the transcriptome, proteome and secretome, impacting pathways related to cell cycle regulation and interferon signaling. Interrogation of multiple independent *L1* elements revealed both shared and distinct cellular responses to acute expression. Thus, our findings demonstrate that acute *L1* expression induces coordinated molecular remodeling extending beyond canonical retrotransposition-associated pathways, rather than simply mimicking senescence or triggering antiviral signaling. This study establishes a comprehensive multi-omics resource for investigating the impact of *L1* transcription in human cells and highlights the need to consider individual transposable element families as distinct regulators of cellular state during aging and disease.

## Introduction

Transposable elements (TEs) are repetitive genetic elements that constitute almost half (∼45%) of the human genome and have long been regarded as “junk DNA” [1–4]. However, it is now well-established that TEs contribute to genome regulation, chromatin organization, and cellular homeostasis across diverse biological contexts and disease [1, 4, 5]. Among these elements, Long INterspersed Element-1 (*LINE-1 or L1*) is an autonomously active retrotransposon in humans, accounting for approximately 17% of the genome and retaining the capacity to mobilize through a copy-and-paste mechanism involving its encoded proteins ORF1p and ORF2p [1, 2, 5].

Like other TEs, *L1* is found strictly repressed under healthy normal conditions and its derepression has been closely linked to aging and age-associated cellular phenotypes [6–8]. Multiple studies have shown that *L1* expression increases with age across species and cell types, including in senescent human fibroblasts [9], aged tissues [10, 11], and age-related diseases [12]. In senescent cells, *L1* derepression has been implicated in the maturation of the senescence-associated secretory phenotype (SASP) and chronic inflammation [9], key hallmarks of aging [13]. In particular, *L1* reverse transcription can generate cytosolic DNA intermediates that are sensed by pattern recognition receptors such as cGAS, leading to activation of the STING pathway and downstream interferon (IFN) signaling [9]. Consistent with this, increased L1 activity has been linked to interferon responses in senescent cells and age-associated inflammation, and pharmacological inhibition of reverse transcriptase has been shown to attenuate inflammatory phenotypes [14]. Furthermore, genetic variants significantly associated with *LINE-1* RNA levels have been found to also be associated with aging markers and phenotypes [15].

These observations have led to the hypothesis that *L1* activation may actively contribute to aging processes [8]. However, the causal relationship between *L1* derepression and aging remains incompletely understood. While studies suggest that *L1*-derived nucleic acids can drive inflammatory signaling through innate immune pathways, it is unclear whether *L1* expression is sufficient to induce broader aging-associated cellular changes or whether it acts as a secondary consequence of aging-related chromatin alterations. Moreover, whether *L1* influences other hallmarks of aging has not been systematically explored.

In a recent study, we showed that acute overexpression of an *Alu* retrotransposon can induce widespread remodeling of the transcriptome, proteome, and secretome, impacting pathways related to mitochondrial function, the cell cycle, and the extracellular matrix [16]. These findings further raise the possibility that TEs can actively regulate cellular physiology rather than acting solely as passive downstream byproducts of age-related failure of genomic surveillance mechanisms. However, whether *L1* exerts similar or distinct effects remains an open question, particularly given its unique capacity to encode functional proteins and mobilize autonomously – in contrast to *Alu*.

Here, we sought to systematically characterize the molecular consequences of *L1* expression on proliferating human primary cells. Using a multi-omics approach encompassing unbiased transcriptomic, proteomic, and secretomic profiling, we analyzed the impact of transient *L1* overexpression across multiple constructs and cell types. These included full-length *L1* elements, constructs expressing individual *L1* open reading frames, and variants with or without reverse transcriptase activity. This design enabled us to begin to disentangle the contributions of *L1* structural components and catalytic activity to downstream cellular phenotypes.

## Results

### Replicative senescence is accompanied by transcriptional derepression of various transposable elements in primary fibroblasts

Studies of cell senescence have previously revealed that it is a highly relevant physiological state for the study of transposon biology [9]. Thus, we first leveraged a publicly available mRNA-sequencing dataset of WI-38 human embryonic fibroblasts undergoing replicative senescence (GSE175533) [17], in order to determine how cellular senescence impacted the landscape of transposable element [TE] expression in primary human fibroblasts. Specifically, we used DESeq2 and TEtranscripts to identify TEs whose RNA levels would change monotonously with increasing population doublings [PDL], starting from PDL 45 (proliferating, high passage), to PDL 52 and PDL 53 (arrested, in deep senescence; **Fig. 1A**). Multidimensional scaling (MDS) of transposon expression levels revealed a clear separation of transcriptomic samples with increasing PDL along the first dimension (**Fig. 1B**), consistent with a robust remodeling of the TE landscape with senescence established in previous studies [9]. Differential expression analysis identified 224 repeat subfamilies exhibiting increased expression and only 17 exhibiting decreased expression with progressive population doublings (**Fig. 1C; Supplementary Table S1A**). Interestingly, observed changes encompass all classes of TEs, including DNA transposons, long interspersed elements (LINEs), short interspersed elements (SINEs), and long terminal repeat (LTR) transposons (**Fig. 1C**).

**Fig. 1.**
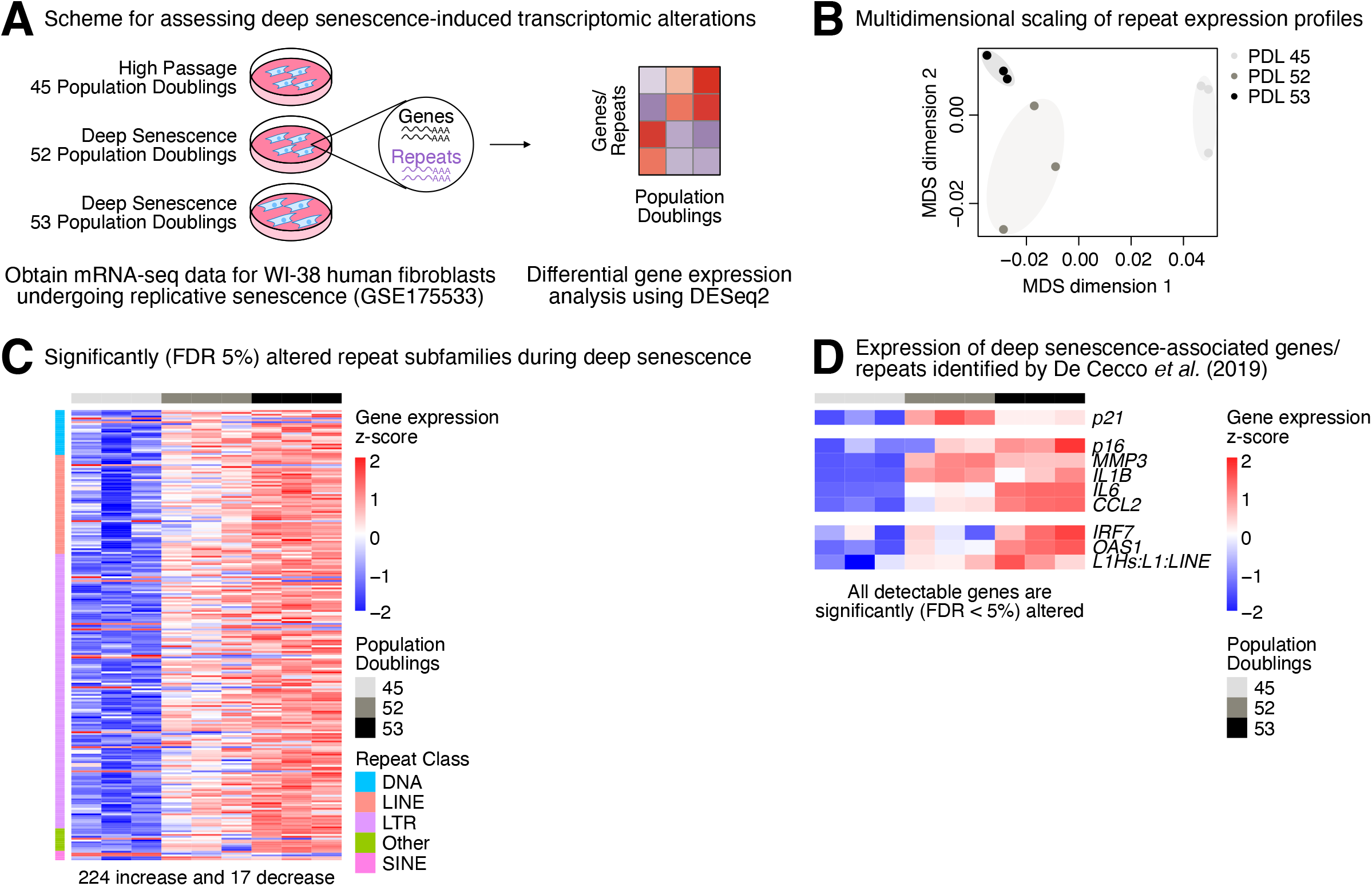
Repetitive elements, including *LINE* retrotransposons, are upregulated in primary human fibroblasts undergoing deep senescence. **(A)** A diagram illustrating the transcriptomic analysis carried out using a publicly available mRNA-sequencing dataset (GSE175533) for primary human fibroblasts of three different population doublings (45, 52 & 53) undergoing replicative senescence. **(B)** Multidimensional scaling (MDS) analysis of repeat element expression profiles. **(C)** Gene expression heatmap for significant (FDR < 0.05) differentially expressed repeat subfamilies across population doublings. **(D)** Expression of deep senescence-associated genes and repeats identified by De Cecco *et al*. 2019 across the three population doublings. FDR: False Discovery Rate.

We next focused on the expression of genes and TEs that are part of a previously identified signature of senescent human embryonic lung LF1 fibroblasts [9], to determine how this signature was regulated in this independent dataset. Consistent with previous work, increased PDL was accompanied by robust upregulation of cell cycle regulator genes (e.g. *CDKN1A*/*p21, CDKN2A/p16*), cytokine genes (e.g. *IL6, IL1B*), interferon-related genes (e.g. IRF7), as well as *L1Hs* (the evolutionarily youngest and only transposition-competent *L1* subfamily in humans; **Fig. 1D**), although the expression of specific interferon genes was below the limit of expression detection in that dataset. Thus, cellular senescence in human fibroblasts is robustly associated with upregulation of a specific transcriptional signature.

### *L1Hs* overexpression induces widespread remodeling of primary human fibroblast multi-omic landscapes

Although *L1* upregulation has been observed in senescent cells [9] (**Fig 1**), it is still unclear whether other features that co-occur with *L1* derepression in deep senescence (*e.g*. cytokine/interferon response, cell cycle arrest, etc.) are mere incidental correlates or whether increased *L1* levels themselves could directly regulate these other pathways, even in non-senescent proliferating cells. Indeed, based on our previous work using a transposition-incompetent *Alu* element [16], we reasoned that overexpressing a transposition-competent *L1* element should have an even stronger influence on cellular phenotypes. For this purpose, we settled on the use of an acute overexpression of a well-established, modified *L1Hs* copy (*LRE3*), which harbors a green fluorescent protein (GFP) reporter in its 3’UTR to track cells with *de novo LRE3* insertions (GFP-LRE3) [18]. Importantly, we first confirmed that this modified *L1* is indeed transposition-competent using transient transfection in HEK293T cells, since they are highly transfectable with high viability (reducing the need for selection to observe the relatively rare occurrence of successful transposition events; **Supplementary Fig. S1A**). Consistent with the overexpressed *L1* being transposition-competent, we observed ∼6-10% GFP^+^ cells 4 days after transfection (**Supplementary Fig. S1B, C; Supplementary Table S1B**).

Next, to test whether *L1* expression could rewire molecular and cellular landscapes of primary, non-senescent fibroblasts, we transiently transfected human IMR-90 fibroblasts using electroporation to overexpress the GFP-LRE3 *L1Hs* element (**Fig. 2A; Supplementary Fig. S2A-C**). To reduce the potential impact of differential cell viability, cells were re-seeded at equal densities after an initial 16h post-transfection recovery period. After reseeding, cells were cultured in low serum media based on recommendations for profiling of secreted proteomes of senescent and/or quiescent cells [19], similar to our previous study [16]. By promoting quiescence, low serum culture conditions control for potential differences in molecular landscapes merely stemming from non-proliferating vs proliferating states. We hypothesized that this might be important for our study since *L1* overexpression in primary fibroblasts may induce senescence-associated changes (**Fig. 1** and De Cecco *et al*. 2019 [9]). After 72h of culturing, cell pellets and corresponding conditioned media were collected for transcriptomic, proteomic, and secretomic analysis (**Fig. 2A**). Importantly, before further profiling, we confirmed expression of the plasmid-specific *LRE3 L1* element using reverse transcription (RT) followed by endpoint polymerase chain reaction (PCR) (**Supplementary Fig. S2A**).

**Fig. 2.**
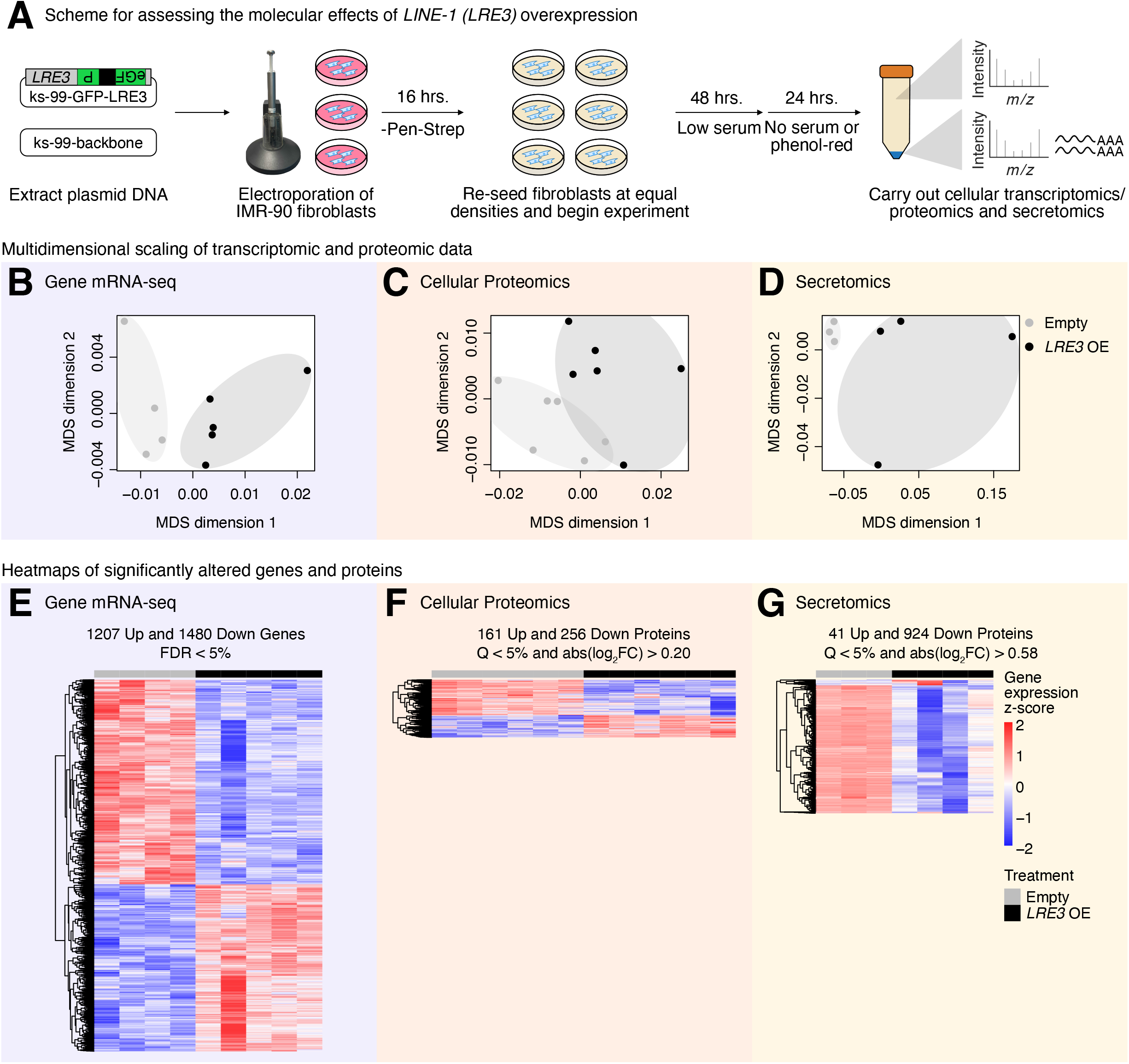
Full-length *L1* overexpression promotes widespread transcriptional and proteomic remodeling in human primary fibroblasts. **(A)** A diagram illustrating how control and full-length *L1* overexpressing IMR-90 fibroblast samples were prepared for multi-omic profiling. N = 6 replicates were independently transfected per group, and N = 4-5 replicates were analyzed by mRNA-sequencing, N = 6 replicates were analyzed by cellular proteomic profiling, and N = 3-4 replicates were analyzed by secretome profiling. Multidimensional scaling (MDS) analysis of the **(B)** genic transcriptome, **(C)** cellular proteome, and **(D)** secretome. Abundance heatmaps for significant **(E)** differentially expressed genes (FDR < 0.05), **(F)** differentially abundant cellular proteins (Q < 5% and abs(log_2_FC) > 0.20), and **(G)** differentially secreted proteins (Q < 5% and abs(log_2_FC) > 0.58). FDR: False Discovery Rate, Q: Q-value.

Consistent with broad cellular effects of *L1* expression in IMR-90 fibroblasts, MDS showed clear separation of control and *L1*-overexpressing cells at all three omics layers: transcriptome (**Fig. 2B**), cellular proteome (**Fig. 2C**), and secretome (**Fig. 2D**). To note, we were also able to detect overexpression of *LRE3* ORF1 in transfected cells using our RNA-seq data (**Supplementary Fig. S2B**). We next asked which specific molecular features may be significantly remodeled in response to *L1* overexpression in primary, non-senescent, IMR-90 human fibroblasts. At the transcriptomic level, we identified 1,207 genes that were significantly upregulated and 1,480 that were significantly downregulated (FDRO<O5%; **Fig. 2E**; **Supplementary Table S1C**). At the level of the cellular proteome, we identified 161 significantly upregulated and 256 significantly downregulated proteins (QO<O5% and abs(log_2_FC)O>O0.20; **Fig. 2F**; **Supplementary Table S1D-F**). Finally, at the level of the secretome, we identified 41 significantly upregulated and 924 significantly downregulated proteins (QO<O5% and abs(log_2_FC)O>O0.58; **Fig. 2G**; **Supplementary Table S1G-H**). Thus, our unbiased multi-omic dataset demonstrates that acute *L1* overexpression is sufficient to drive broad molecular reprogramming across ‘omic’ layers in primary human fibroblasts.

### *L1Hs* overexpression leads to changes in cell proliferation and inflammatory-related pathways

Next, we asked how observed molecular changes in *L1*-overexpressing cells could drive functional remodeling, by leveraging Gene Set Enrichment Analysis (GSEA) [20] for each individual “omic” layer, using gene set collections from Reactome [21] and MSigDB Hallmarks [22] (**Fig. 3A**; see **methods**). Importantly, GSEA using Reactome gene set definitions revealed molecular alterations dominated by upregulation of cell cycle–related processes at the transcriptomic (**Fig. 3B**), proteomic (**Fig. 3C**) and secretomic (**Fig. 3D**) levels. Specifically, at the transcriptome level, top enriched gene sets included “cell cycle, mitotic”, “cell cycle checkpoints”, “M phase”, “mitotic prometaphase”, and “resolution of sister chromatid cohesion” (**Fig. 3B**; **Supplementary Table S2A**). These findings were consistent at the cellular proteome level, with strong upregulation of proteins related to “Mitotic G1 phase and G1/S transition” and “cell cycle, mitotic” (**Fig. 3C**; **Supplementary Table S2B**). The upregulation of cell cycle-related processes was also clear at the secretome level, with enriched terms including “condensation of prophase chromosomes” and “meiotic recombination” (**Fig. 3D**; **Supplementary Table S2C**). Importantly, we also observed downregulation of genes/proteins related to extracellular matrix [ECM] biology, including “extracellular matrix organization” and “degradation of the extracellular matrix” (**Fig. 3B,D**; **Supplementary Table S2A,C**). ECM-related terms were also significantly impacted in response to *Alu* overexpression in our previous work [16], albeit in the opposite direction, suggesting that cellular responses to TE expression are likely to recurrently lead to ECM remodeling, consistent with known age-related fibrosis and tissue stiffening [23]. Importantly, analyses using the MSigDB Hallmark gene sets were largely consistent with Reactome findings (**Fig. 3E-G**; **Supplementary Table S2D-E**), revealing upregulation of cell cycle-related terms (*e.g*. “G2M checkpoint”, “mitotic spindle”, “E2F targets”) and downregulation of ECM related terms (*e.g*. “epithelial mesenchymal transition”). To note, the MSigDB Hallmark analysis also revealed significant misregulation of DNA damage and repair pathways (*e.g*. “DNA repair”, “UV response down”), consistent with the documented genotoxic impact of transposition-competent *L1* (**Fig. 3E, G**). To our surprise, gene sets classically related to SASP and inflammation (*e.g*. Reactome “signaling by interleukins” and “interferon gamma signaling” and MSidgDB “interferon gamma response” and “inflammatory response”) were significantly downregulated upon *L1* overexpression in the transcriptomic layer (**Fig. 3B,E**; **Supplementary Table S2A**), suggesting that cellular immune responses may be suppressed – rather than activated – in this context. Although these transcriptome-level changes may be adaptive, downstream responses after 72h, our observations suggest that cellular responses to L1, including through innate immunity pattern recognition receptors, may be context-specific (see more below).

**Fig. 3.**
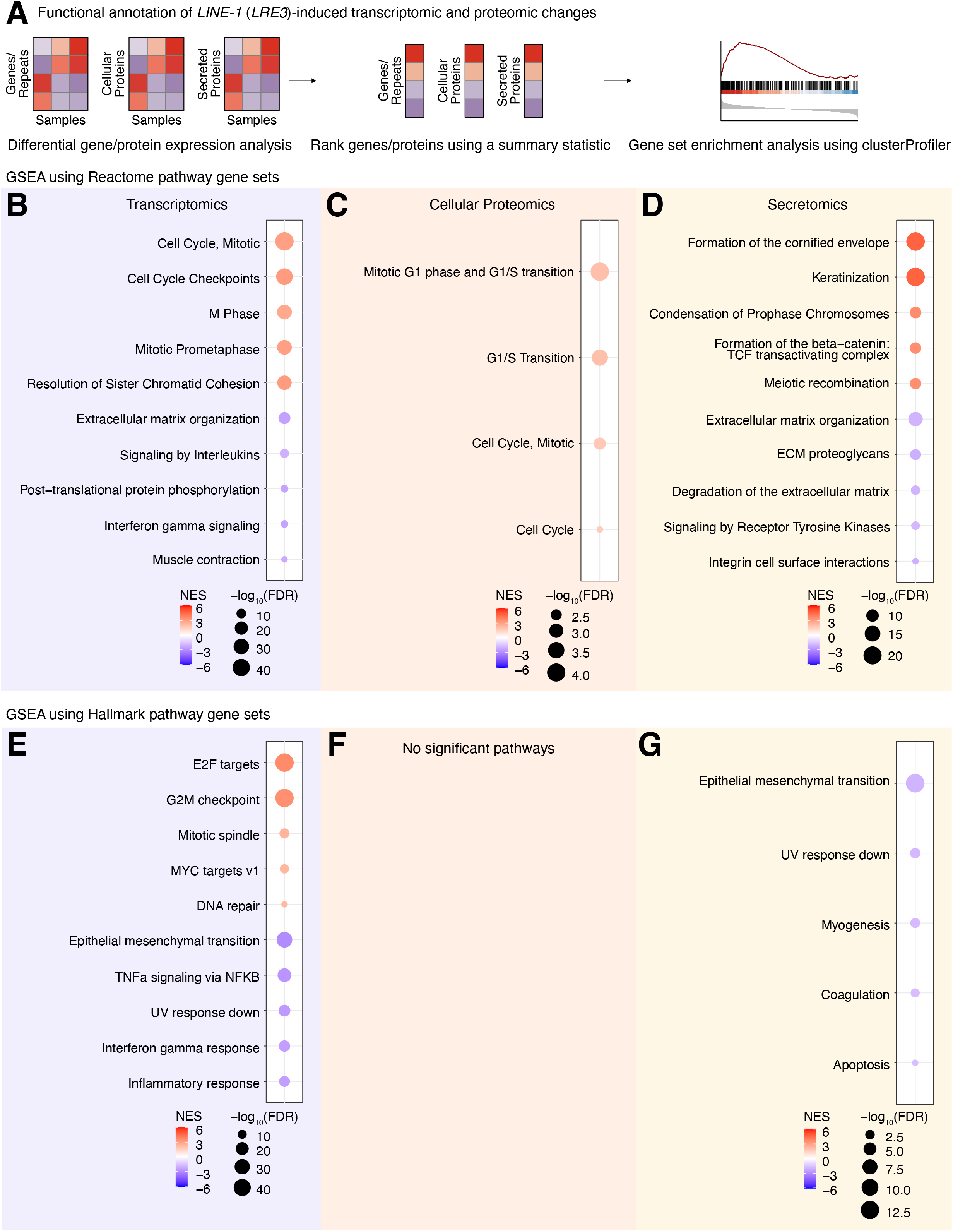
Full-length *L1* overexpression drives molecular changes in cell cycle and interferon-related signaling pathways in human primary fibroblasts. **(A)** A diagram illustrating how functional annotations were assigned to transcriptomic and proteomic changes with gene set enrichment analysis. The top 5 significant (FDR < 0.05) Reactome pathway gene sets in each direction for the **(B)** transcriptome, **(C)** cellular proteome, and **(D)** secretome were plotted. The top 5 significant (FDR < 0.05) Hallmark pathway gene sets in each direction for the **(E)** transcriptome, **(F)** cellular proteome, and **(G)** secretome were also plotted. FDR: False Discovery Rate, NES: Normalized Enrichment Score.

Next, we decided to functionally validate whether functional enrichments related to cell cycle were related to actual changes in cell proliferation. For this purpose, we performed a propidium iodide-based cell cycle analysis (**Fig. 4A**; **Supplementary Fig. S4A**; **Supplementary Table S3A**). As predicted by our multi-omic analysis, acute *L1* overexpression led to cells exhibiting a modest, non-significant decreased trend in the proportion of cells in G0/G1 phase (p ∼0.06) and a significant accumulation of cells in S phase (p < 0.05; **Fig. 4B**). Most likely, these findings are compatible with the notion that acute L1 expression leads to cell cycle arrest at the DNA replication checkpoint. To note, these findings are reminiscent of our previous observations in response to acute expression of an old, transposition-incompetent *Alu* element in IMR-90 cells [16], suggesting that disruption of cell cycle-related programs may be a common response to TE upregulation in primary human fibroblasts.

**Fig. 4.**
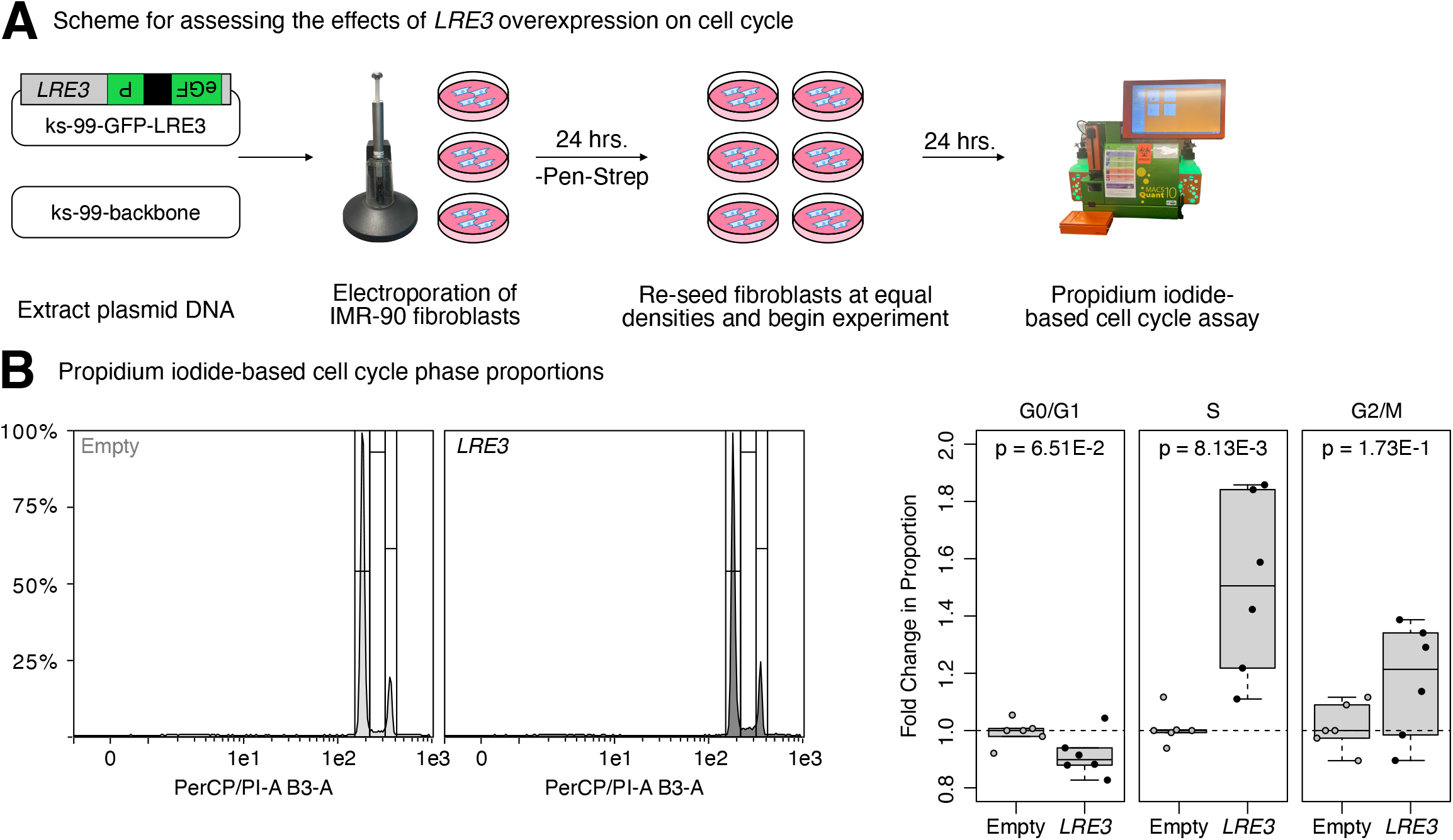
Full-length *L1* transient overexpression drives cell cycle alterations in primary human fibroblasts. **(A)** A diagram illustrating how the functional effects of *L1* overexpression on the cell cycle were assessed. **(B)** Representative flow cytometry histograms for propidium iodide-stained control and *L1* overexpressing cells. Two experiments were combined, for a total of N = 6 per group, and the relative number of cells in G0/G1, S, and G2/M phases in the control and overexpression group were compared with a Wilcoxon rank sum test. p < 0.05 was considered significant.

Since many of the processes regulated in response to *L1* overexpression were related to hallmarks of aging [13], we next carried out GSEA using curated aging- and senescence-related gene sets (**Supplementary Fig. S3A**). Interestingly, we observed a negative relationship between the transcriptional response to *L1* overexpression and the responses upregulated and downregulated with aging across human tissues from the Genotype-Tissue Expression (GTEx) project [24], deep replicative senescence in fibroblasts (**Figure 1**), and general cell senescence according to CellAge [25] (**Supplementary Fig. S3B; Supplementary Table S2F-H**). Intriguingly, we observed significant downregulation of senescence-associated genes sets, including SenMayo [26] and the core SASP [27] (**Supplementary Fig. S3B**). Consistent with this, we also observed a downregulation of individual gene markers previously linked to deep senescence and L1 upregulation in human fibroblasts [9] (**Supplementary Fig. S3C**). As an important control, the directionality of regulation for the same gene set collection in our re-analysis of the deep senescence transcriptome (**Figure 1**) revealed regulation mostly consistent with accelerated aging (with the notable exception of the core SASP gene set), as expected based on the reported accumulation of senescent cells with aging (**Supplementary Fig. S3D; Supplementary Table S2I**). Together, our analyses support the notion that acute *L1* overexpression in proliferating fibroblasts does not merely initiate senescence or senescence-like molecular remodeling.

Collectively, our analyses support the notion that *L1* overexpression drives a coordinated reprogramming of human fibroblast multi-omic landscapes, including induction of pathways related to cell proliferation and suppression of genes related to inflammatory signaling. Interestingly, a similar biological antagonism, where interferon gamma was substantially reduced during the S-G2/M phases of the cell cycle, was previously reported in macrophages in mice [28].

### Full-length *L1* expression in non-senescing, non-cancerous cells suppresses interferon signaling

Next, we decided to further explore our surprising observation of decreased inflammatory and interferon signaling in response to *L1* overexpression in IMR-90 primary fibroblasts, which was observed at all ‘omic’ levels (**Fig. 3**; **Supplementary Fig. S5A,B; Supplementary Table S2J-O**). This finding was especially surprising to us, as the general consensus in the field is that, due to their viral origin, TEs in general, and *L1* in particular, are expected to lead to activation of anti-viral pathways like interferon signaling downstream of recognition by pattern recognition receptor pathways cGAS/STING or MDA5/RIG-I/MAVS [9, 29–31].

To help resolve our puzzling observation in this matter, we thus decided to systematically test the impact on interferon signaling pathways from (i) different full-length *L1* constructs (i.e. *L1* under its native promoter with and without a retrotransposition reporter cassette [*LRE3*, *L1RP*], CMV-driven *L1* [CMV-*L1RP*], codon-optimized *L1* [*ORFeus*]), (ii) different cellular proliferation states (*i.e*. quiescent *vs*. proliferating cells), (iii) different fibroblast isolates (*i.e*. IMR-90 vs. WI-38 primary human fibroblasts), and (iv) different cell transformation/immortalization states (*i.e*. primary [IMR-90, WI-38], immortalized [HEK293T, RPE] and cancerous [HeLa] cells) (**Fig. 5A-B**; **Supplementary Fig. S6-12; Supplementary Table S4A-AN**). In addition, we reanalyzed a publicly available mRNA-sequencing dataset for telomerase-immortalized retinal pigment epithelium-1 (RPE) cells harboring a doxycycline-inducible codon-optimized *L1* (*ORFeus*) or luciferase control (GSE119999; **Supplementary Fig. S13A; Supplementary Table S4M, Z, AM**) [32]. Specific constructs, experimental design and related dataset validation are reported in **Supplementary Table S4A**.

**Fig. 5.**
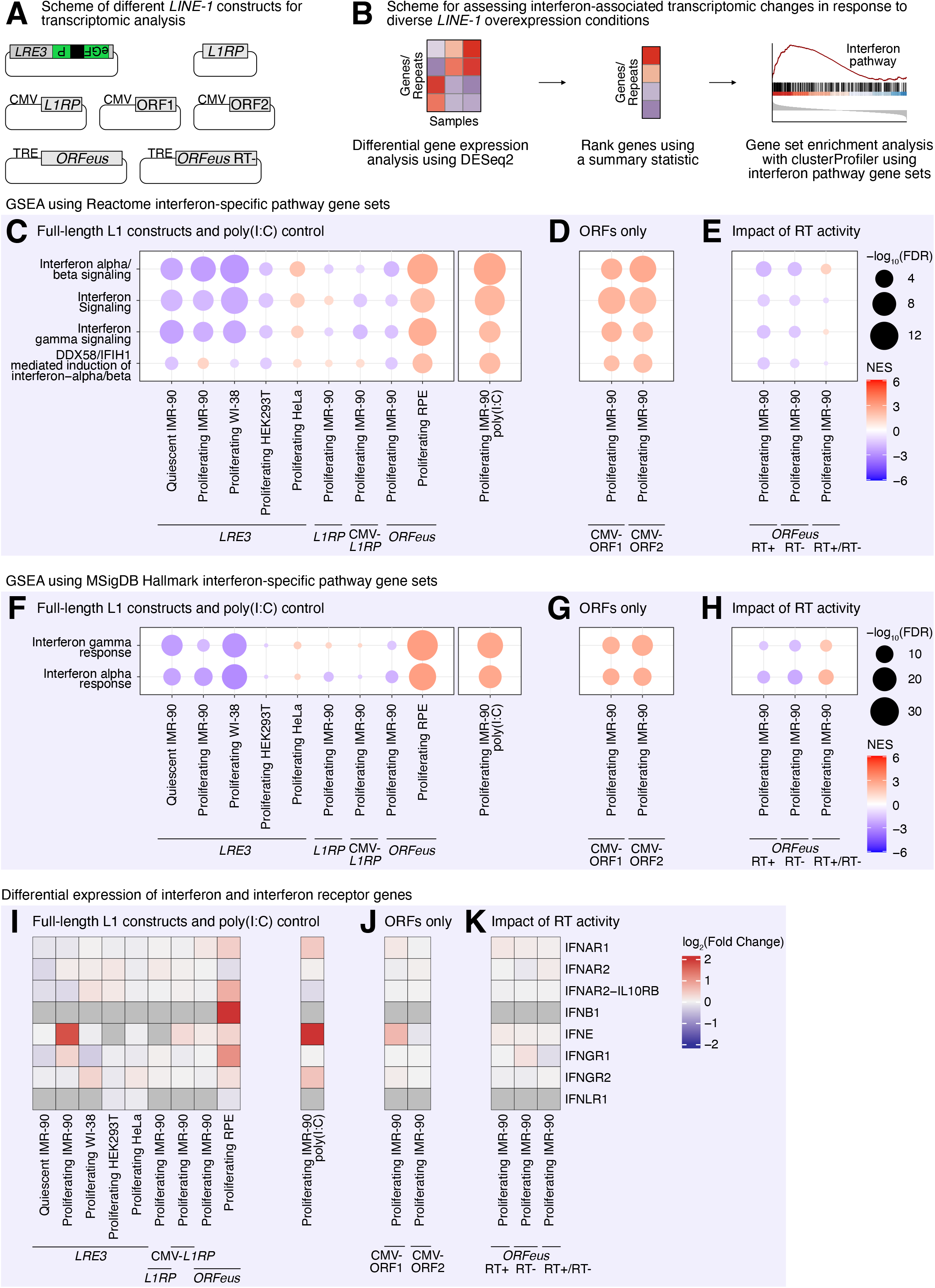
Context-dependent regulation of the interferon signaling pathway in response to *L1* overexpression. **(A)** An illustration of the constructs used to assess the transcriptomic impact of *L1* overexpression. **(B)** A diagram illustrating how interferon-associated transcriptomic changes in response to diverse *L1* constructs in diverse cell types were assed. GSEA analysis using Reactome interferon-specific pathway gene sets for diverse cell types overexpressing **(C)** full-length *L1* constructs, **(D)** only *L1* ORFs, and **(E)** *L1 ORFeus* constructs with functional and non-functional reverse transcriptase (RT). GSEA using MSigDB Hallmark interferon-specific pathway gene sets for diverse cell types overexpressing **(F)** full-length *L1* constructs, **(G)** only *L1* ORFs, and **(H)** *L1 ORFeus* constructs with functional and non-functional reverse transcriptase (RT). Differential expression of interferon and interferon receptor genes in diverse cell types overexpressing **(I)** full-length *L1* constructs, **(J)** only *L1* ORFs, and **(K)** *L1 ORFeus* constructs with functional and non-functional reverse transcriptase (RT). FDR: False Discovery Rate, NES: Normalized Enrichment Score.

Consistent with our results using the *LRE3* (*L1Hs*) reporter construct in quiescent IMR-90 cells (**Fig. 3**; **Supplementary Fig. S5A,B**), we generally observed downregulation of interferon-related gene sets in response to full-length *L1* constructs (*LRE3, L1RP, ORFeus*) in non-cancerous cells (IMR-90, WI-38, HEK-293T), regardless of proliferation status (**Fig. 5C**, **5F**). In contrast, cancerous cells (HeLa, RPE) showed the expected upregulation of interferon-related gene sets (**Fig. 5C**, **5F**). Although transcriptional responses to the *L1RP* construct in IMR-90 fibroblasts were more nuanced than to *LRE3*, significant downregulation of select interferon-related pathways was still observed, and strong downregulation of interferon related pathways was observed in response to the distinct *ORFeus* construct (**Fig. 5C**, **5F**). Thus, our systematic analyses suggest that downregulation of interferon-related genes in *LRE3*-transfected IMR-90 cells is a robust result, and not a mere byproduct of the specific construct (GFP-*LRE3*) or fibroblast cell line (IMR-90) chosen for our unbiased multi-omic resource. Importantly, we confirmed that, using the same gene set definitions, IMR-90 human primary fibroblasts can transcriptionally upregulate interferon-related responses in response to poly(I:C) exposure (a viral infection mimic; **Fig. 5C**, **5F; Supplementary Fig. S13B-C; Supplementary Table S4N, AA, AN**), supporting the hypothesis that our surprising observations are likely to be a meaningful response to acute, full-length *L1* expression in a previously untested cellular context (primary, non-senescent fibroblasts).

Intriguingly, transcriptional suppression of interferon-related pathways was no longer observed when *L1RP* ORF1 and ORF2 were expressed independently from each other under a CMV promoter (**Fig. 5D**, **5G**), suggesting that cellular responses to *L1* vary with the partial or full-length transcripts being overexpressed. Indeed, *L1* can produce several truncated transcripts due to the presence of internal canonical and noncanonical polyadenylation signals [33], and elevated levels of truncated ORF1p can suppress *L1* mobilization *in trans* [34], potentially altering downstream interferon signaling. Additionally, an *L1* ORF1 RNA-binding mutant was shown to promote the expression of interferon-alpha [30], suggesting that alterations in ORF1 RNA binding partially mediate interferon signaling; this binding may be perturbed following ORF1-specific overexpression. Finally, high levels of ORF1p have been associated with either higher or lower interferon signaling in different cancer types [35], highlighting the multifactorial nature of *L1*-associated interferon signaling. Similarly, truncated ORF2 proteins have been shown to retain the potential to cause toxicity and DNA damage even in the absence of the rest of the transcript [36]; part of the toxicity may be due to reverse transcriptase [RT]-generated, immunostimulatory cDNA products. As such, we assessed whether the RT activity of ORF2, required for *L1* transposition, played any role in the observed transcriptional suppression of interferon-related pathways in IMR-90 fibroblasts, using previously published *ORFeus*-derived constructs [32]. Although both RT^+^ and RT^−^ versions of the *ORFeus L1* led to interferon-related pathway downregulation in IMR-90 fibroblasts (**Fig. 5E**, **5H**), RT^+^ *ORFeus*-expressing IMR-90 cells had significantly higher activation of interferon-related pathways than RT^−^ *ORFeus*-expressing IMR-90 cells (RT^+^/RT^−^ column; **Fig. 5E**, **5H**), meaning that interferon-related pathway downregulation was weaker with uncompromised RT activity.

Given these observations, we next examined the expression of specific interferon-related genes across *L1* overexpressing cell datasets. Largely, interferon and interferon receptor genes were not differentially expressed across *L1* overexpressing constructs relative to controls (**Fig. 5I-K; Supplementary Fig. S14A**), suggesting that either (i) the negative enrichment of interferon-related pathways could be driven by subtle, coordinated changes in a subset of genes within these gene sets rather than large-scale suppression of canonical interferon genes or (ii) the enrichment is related to changes in gene expression for genes upstream or downstream of the interferons and their direct receptors. Thus, our observations are consistent with the notion that competing signals are activated in response to full-length *L1* expression in primary human fibroblasts, including signals that have divergent, contradicting impacts on interferon signaling responses.

### Multi-contrast functional enrichment analysis identifies robust regulation of processes related to the hallmarks of aging

Next, we refocused on our multi-omic *LRE3*-transfected IMR-90 quiescent primary fibroblast dataset, in order to better pinpoint robust alterations in response to *L1* overexpression across omic layers (**Fig. 6A**). We first performed a basic overlap analysis of significantly regulated genes/corresponding proteins across transcriptomic, cellular proteomic, and secretomic datasets (**Fig. 6B**; **Supplementary Table S5A**). This analysis led us to identify 59 genes/corresponding proteins consistently altered across layers (**Fig. 6B**). Interestingly, when focusing on these specific gene products transcriptomic levels and intracellular proteome showed the strongest correlation (Spearman ρ = 0.63) (**Supplementary Table S5B)**. To note, 16 of these genes/proteins were associated with ECM organization, including seven collagen genes (*COL1A1, COL1A2, COL3A1, COL5A2, COL6A1, COL6A3*, and *COL14A1*) [37], consistent with strong remodeling of ECM-related pathways gene regulation. In addition, *TGFB1*, a key regulator of cell growth and tissue repair [38], and *HMGB2*, a chromatin associated gene [39], were also consistently altered across layers (**Supplementary Table S5A**).

**Fig. 6.**
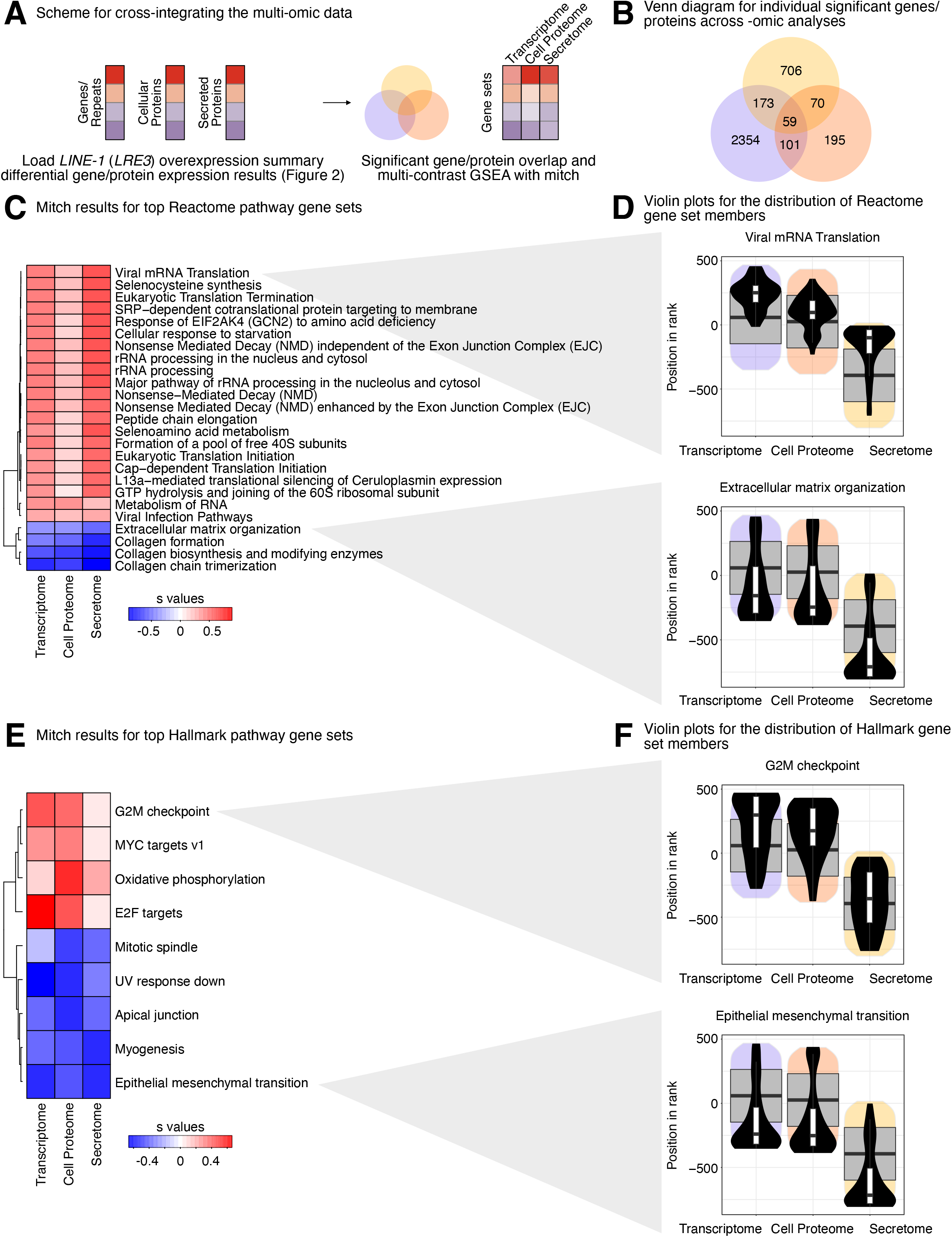
Integrated multi-omic functional enrichment analysis following L1 overexpression in primary human fibroblasts highlights alterations in viral mRNA translation, extracellular matrix organization, and cell cycle processes. **(A)** A diagram illustrating how transcriptome, proteome, and secretome changes were integrated via overlap analysis or multi-contrast gene set enrichment analysis. **(B)** Venn diagram comparing individual genes and proteins that significantly changed in each “-omics” analysis. The top multi-contrast gene set enrichment results using **(C)** Reactome pathway gene sets and **(E)** Hallmark pathway gene sets. Gene sets with an adjusted MANOVA p-value < 0.05 were considered significant. Violin plots for select **(D)** Reactome pathway gene sets and **(F)** Hallmark pathway gene sets related to recurring pathway themes are shown.

Gene-level multi-level ‘omic’ integration tends to be underpowered. Thus, for increased sensitivity, we next performed multi-contrast functional enrichment analysis across ‘omic’ layers by leveraging the rank-based MANOVA framework implemented by ‘mitch’ [40], an approach designed to extract robust conclusions from multi-omic studies (**Fig. 6C-F; Supplementary Table S5C-D**). Mitch analysis revealed strong, directional agreement across ‘omic’ layers for *LRE3*-transfected IMR-90 quiescent primary fibroblasts (**Fig. 6C,E**). Interestingly, we observed strong multi-omic evidence for upregulation of protein homeostasis-related gene sets (e.g. related to ribosome function and rRNA metabolism), including Reactome “eukaryotic translation initiation”, “peptide-chain elongation”, and “rRNA processing” (**Fig. 6C**). Consistent with the virus-related nature of *L1* sequences, there was multi-level support for upregulation of viral biology-related pathways, including Reactome “viral mRNA translation” and “viral infection pathways” (**Fig. 6C,D; upper panel**). Consistent with our ‘omic’ layer-specific analyses, we observed strong evidence for downregulation of ECM-related processes, most dramatically in the secretome layer, including Reactome “extracellular matrix organization” and “collagen biosynthesis and modifying enzymes” (**Fig. 6C,D; lower panel**) and MSigDB Hallmarks “epithelial mesenchymal transition” (**Fig. 6E,F; lower panel**). We also observed consistent upregulation of processes related to nutrient sensing, notably Reactome “cellular response to starvation” (**Fig. 6C**). In addition, there was strong multi-omic signal for upregulation of cell cycle-related genes, consistent with our single ‘omic’ layer analyses, including MSigDB Hallmark “G2/M checkpoint” and “E2F targets” (**Fig. 6E, F; upper panel**). Finally, we also observed robust multi-level evidence for deregulation of DNA-damage sensing/repair pathways, specifically MSigDB Hallmark “UV response down” (**Fig. 6E**). Thus, integrated analysis of *LRE3*-transfected IMR-90 quiescent primary fibroblasts reveals strong multi-omic regulation of pathways related to the hallmarks of aging.

Next, we used a similar approach of multi-contrast functional enrichment analysis using the ‘mitch’ framework, this time leveraging our various transcriptomic datasets of full-length *L1* construct overexpression in IMR-90 fibroblasts (**Fig. 7A; Supplementary Table S5E-F**). Indeed, we reasoned that this approach would allow us to zoom in on *L1*-driven responses, and help filter out potential noise related to *L1* construct-specific idiosyncrasies. Interestingly, similar to our interferon-related gene specific analyses, we saw much stronger agreement between *LRE3* and *ORFeus* expression conditions, with more divergence compared to the *L1RP* conditions (**Fig. 7B,D**). To note, this orthogonal approach picked up a robust downregulation of interferon-related signaling in response to *L1* overexpression (*e.g*. MSigDB Hallmark “interferon gamma response” and “interferon alpha response”; **Fig. 7D**), consistent with our earlier hypothesis-driven analysis (**Fig. 5**; **Supplementary Fig. S5**).

**Fig. 7.**
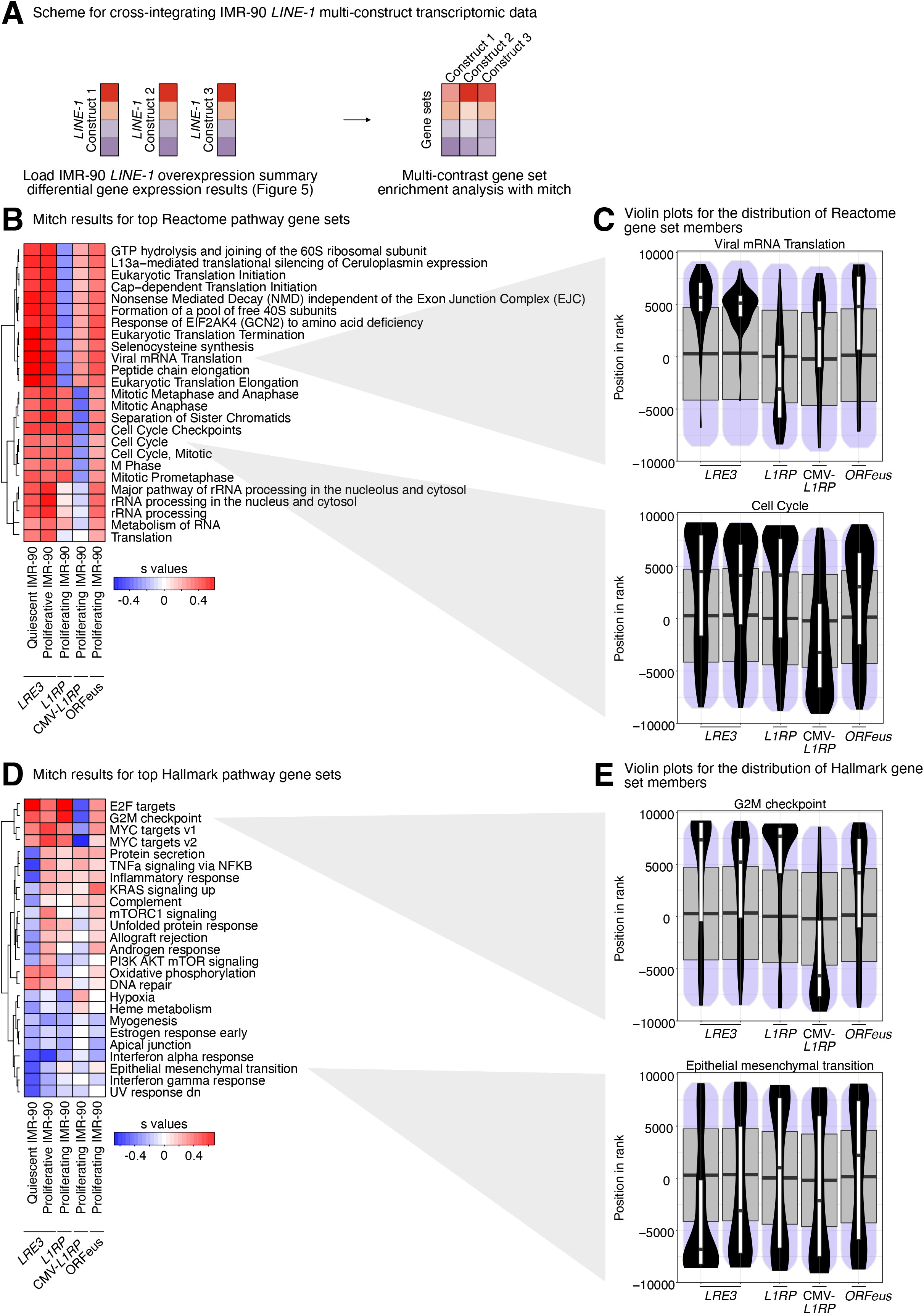
Multi-contrast integration of L1 element overexpression highlights alterations in cell cycle, viral mRNA translation, and proteostasis. **(A)** A diagram illustrating how transcriptomic changes resulting from different full-length *L1* constructs were integrated via multi-contrast gene set enrichment analysis. The top multi-contrast gene set enrichment results using **(B)** Reactome pathway gene sets and **(D)** Hallmark pathway gene sets. Gene sets with an adjusted MANOVA p-value < 0.05 were considered significant. Violin plots for select **(C)** Reactome pathway gene sets and **(E)** Hallmark pathway gene sets related to recurring pathway themes are shown.

In line with our *LRE3*/IMR-90-focused multi-omic analysis above (**Fig. 6**), multi-construct multi-contrast analysis across L1 overexpression paradigms revealed robust, largely consistent regulation of gene sets related to protein homeostasis (*e.g*. Reactome “eukaryotic translation initiation”, “peptide-chain elongation”, “rRNA processing”), viral biology/infection (*e.g*. Reactome “viral mRNA translation”), ECM-related processes (*e.g*. MSigDB Hallmark “epithelial mesenchymal transition”), nutrient-sensing (*e.g*. MSigDB Hallmark “PI3K AKT mTOR signaling” and “Reactome response of EIF2AK4 (GCN2) to amino acid deficiency”), cell cycle (*e.g*. MSigDB Hallmark “G2/M checkpoint” and “E2F targets”) and DNA-damage sensing/repair pathways (*e.g*. MSigDB Hallmark “UV response down”) (**Fig. 7B-E**). Together, our data shows that pathways related to the hallmarks of aging are transcriptionally remodeled in response *L1* overexpression in primary fibroblasts.

## Discussion

### A resource to study the impact of *L1* transcription in primary, non-senescent human fibroblasts

Beyond the biological findings, this study establishes a comprehensive multi-omics resource for investigating the consequences of *L1* transcription in primary human fibroblasts. By integrating transcriptomic, cellular proteomic, and secretomic profiling following transient *L1* overexpression and systematically comparing multiple *L1* expression architectures, including (i) full-length constructs, (ii) isolated ORF expression, and (iii) reverse transcriptase functional and non-functional variants, this dataset provides a framework for dissecting how *L1* expression influences cellular physiology. The inclusion of multiple expression systems and independent datasets also permits comparison of conserved and context-dependent responses across primary fibroblasts and transformed cell lines. Rather than examining a single downstream pathway, the resource captures coordinated changes across multiple molecular layers, enabling both pathway-level and gene-level interrogation of *L1*-mediated responses. This represents a significant advance over many previous studies, which typically rely on a single construct or experimental model.

The value of this dataset extends beyond the conclusions presented here. It provides a resource for exploring how *L1* transcription influences diverse biological processes. Because transcriptomic, intracellular proteomic and secretomic measurements were generated from matched experimental systems, the data also enable future studies of post-transcriptional regulation, protein secretion dynamics, and pathway concordance across molecular layers.

### Acute *L1* upregulation in primary, non-senescent human fibroblasts and interferon-related signaling

*L1* derepression is a feature of senescence and, once expressed, *L1* particles can generate ligands that can be recognized by pattern recognition receptors, leading to chronic inflammation [9, 29–31]. A central mechanism is accumulation of cytoplasmic *L1* cDNA, which activates cGAS-STING and induces type I interferon in senescent cells [9]. In a complementary process independent of *L1* cDNA, RNA sensing mechanisms can also drive innate immune signaling by triggering interferon signaling [29–31]. Intriguingly, our findings suggest that the relationship between *L1* expression and interferon signaling is more context-dependent than previously assumed. While endogenous *L1* derepression during aging and senescence is widely associated with innate immune activation, our analyses of acute *L1* overexpression paints a more nuanced picture. Full-length *L1* expression consistently resulted in transcriptional downregulation of interferon-related pathways, whereas isolated expression of ORF1 or ORF2 activated these same pathways, indicating that the cellular response could very well depend on the architecture of the expressed *L1* transcript (more varied when derived from potentially truncated copies in the nuclear genome). Furthermore, although reverse transcriptase activity modestly enhanced interferon pathway activation, both functional, wildtype RT and non-functional, mutant RT ORFeus constructs retained an overall suppressive transcriptional signature, suggesting that reverse transcription contributes to, but does not always drive, the interferon response. The absence of widespread differential expression of canonical interferon genes further indicates that these pathway changes arise from coordinated modulation of downstream pathway components rather than global activation or repression of interferon signaling.

The downstream immune phenotype resulting from *L1* expression is perhaps different depending on cell state. *IFN-I* response has been described as a phenotype of late senescence [9]. Indeed, *PAX5*, an upstream transcriptional regulator of cGAS–STING, showed enriched binding at *L1* 5’-UTRs in senescent cells [41]. In addition, loss of cDNA degraders such as *TREX1* also contributes to constitutive cGAS–STING activation [29]. Hence, this might suggest that *L1* surveillance and nucleic acid clearance might be weakened in senescence leading to sustained activation of the interferon pathway.

### Context-dependency of the molecular cellular responses to *L1* upregulation

An important observation emerging from our study is that the transcriptional response to acute *L1* expression is dependent on cell type and/or state. Full-length *L1* constructs suppressed expression of interferon-related pathways in primary fibroblasts (IMR-90 and WI-38), whereas this effect was attenuated in HEK293T cells and absent or even reversed in HeLa cells. These findings suggest that the consequences of *L1* expression are not intrinsic to the retrotransposon itself but are determined by the regulatory context of the host cell.

Primary fibroblasts likely represent a physiologically-relevant cell model because they retain intact cell-cycle checkpoints, chromatin regulation, and innate immune pathways that are frequently altered during immortalization or oncogenic transformation. In contrast, due to the immortalization and/or oncogenesis process, HEK293T and HeLa cells are likely to host genetic and epigenetic alterations affecting antiviral signaling, DNA damage responses, and cell-cycle regulation. Consequently, transformed cells may not interpret *L1* expression in the same manner as primary cells, explaining why the magnitude and the direction of interferon pathway regulation differed across the cell types examined in this study. Our observations are particularly relevant in contextualizing our results with the existing literature. Much of our current understanding of *L1*-mediated innate immune activation originates from studies performed in senescent cells [9], transformed cell lines [29, 30, 32], or cancer models, where endogenous *L1* derepression has been linked to activation of type I interferon signaling. To note, it is likely that cell senescence, with its accompanying broad changes in the nuclear epigenome, cellular energetics and gene regulation, also represents a unique, distinct context for acute L1 response, which will be interesting to evaluate in future studies (outside of the scope of the present work).

This perspective also provides a framework for reconciling seemingly contradictory observations across the literature. Studies of endogenous *L1* derepression during aging and senescence describe a chronic state in which persistent accumulation of *L1*-derived nucleic acids contributes to sustained innate immune activation and inflammaging. By contrast, the present work examines the acute consequences of experimentally defined *L1* expression paradigms and reveals that these can initiate broader transcriptional reprogramming in which inflammatory and proliferative pathways are simultaneously regulated.

### *L1* upregulation in primary, non-senescent fibroblasts leads to broad, multi-omic regulation of processes related to the hallmarks of aging

In our recent multi-omics analysis of *AluJb* overexpression in primary human fibroblasts [16], *Alu* expression induced widespread remodeling of aging-associated pathways, including repression of cell-cycle, mitochondrial and proteostasis pathways together with activation of extracellular matrix organization and senescence-associated secretory pathways, resembling transcriptional changes observed during physiological aging and cellular senescence. In contrast, although acute *L1* overexpression similarly perturbed cell-cycle progression, ‘omic’ responses were largely distinct from both *Alu* overexpression and natural aging. Rather than repressing cell-cycle programs, *L1* overexpression consistently enriched cell-cycle, *E2F* and *G2/M* checkpoint pathways across transcriptomic and proteomic layers, while extracellular matrix organization and interferon signaling were transcriptionally suppressed. Further, comparison with curated aging and senescence signatures revealed an overall inverse relationship between the *L1* transcriptional response and organismal aging, deep replicative senescence, CellAge, SenMayo and core SASP gene sets. Importantly, the same analytical framework correctly identified positive concordance between the deep senescence reference dataset and markers of both organismal aging and senescence, indicating that the observed inverse relationship with *L1* was not an artifact of the analysis but instead reflected a biologically distinct response to acute *L1* expression.

These observations suggest that different TE families are unlikely to contribute equally to aging biology. Rather than acting as interchangeable markers of genomic dysregulation, individual TEs may regulate distinct aspects of the aging phenotype. *Alu* expression appears to promote a transcriptional program closely aligned with senescence-associated remodeling, whereas *L1* expression induces a broader cellular reprogramming characterized by activation of proliferation-associated pathways and attenuation of inflammatory signaling. Although both elements ultimately perturb cell-cycle homeostasis, the downstream molecular programs they engage differ substantially, implying that they act through distinct regulatory mechanisms.

This distinction has important implications for understanding transposable element biology during aging. Aging and senescence are accompanied by derepression of multiple transposable element families, including *L1*, *Alu* and endogenous retroviruses (ERVs), yet these elements differ markedly in evolutionary origin, coding capacity, RNA structure and mechanisms of host interaction. Consequently, the biological consequences of transposable element derepression are likely to depend not only on the extent of derepression but also on which transposable elements become expressed, in which cell types, and at what stage of the aging process.

Finally, the dataset highlights that *L1* biology cannot be interpreted solely through the lens of retrotransposition or innate immune activation. Instead, it provides evidence that the consequences of *L1* expression depend on cellular context, and the balance of competing transcriptional programs. Our findings also highlight the idea that transposable elements should be viewed as functionally diverse regulators of cellular state rather than a homogeneous class of repetitive elements. As additional studies emerge investigating endogenous *L1* activation during aging, senescence, development, and disease, this resource offers an experimentally controlled reference against which those physiological states can be compared.

Taken together, our molecular multi-omics, -cell, and -L1 construct analyses reveal L1 overexpression-induced changes in pathways related to several hallmarks of aging (**Fig. 8A**). These include changes associated with primary hallmarks (e.g. loss of proteostasis, genomic instability), antagonistic hallmarks (e.g. mitochondrial dysfunction), and integrative hallmarks (e.g. extracellular matrix changes, altered intercellular communication, and cell cycle changes that can manifest as stem cell exhaustion).

**Fig. 8.**
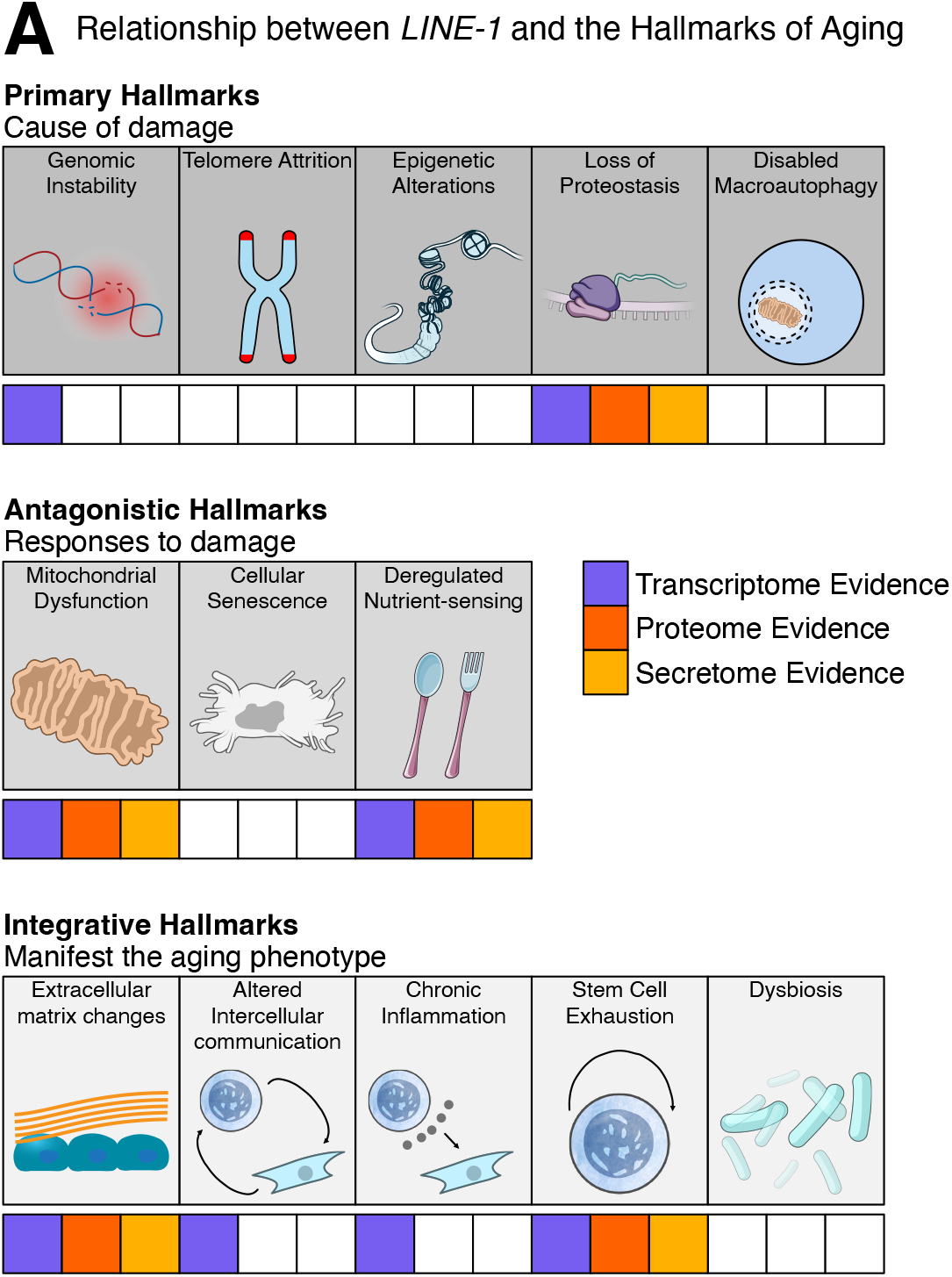
*L1*-induced alterations relate to multiple hallmarks of aging. **(A)** A diagram summarizing the findings from this study as they relate to the hallmarks of aging.

### Limitations of the study

Although we tested several acute *L1* overexpression paradigms, we note that the mechanisms by which *L1* can impact host physiology are multifactorial and not yet completely disentangled. As previously noted, *L1* can produce several truncated transcripts due to the presence of internal canonical and noncanonical polyadenylation signals [33]. As such, the product of *L1* transcription may be a pool of different transcripts whose composition may change in response to different cellular cues, and the composition of this pool may partially dictate the impact of *L1* expression on host cells. Future studies should incorporate long-read sequencing technologies to define these transcripts across experimental conditions, and the use of codon-optimized sequences may help isolate the effects of specific protein isoforms (although this approach comes with its own caveats). Second, both *L1* RNA [29, 30] and its products [9, 14] are believed to perturb host physiology, though it remains unclear which perturbations are RNA-mediated, protein-mediated, or shared. Future studies could incorporate additional *L1* ORF protein mutants, as well as constructs with mutations that disrupt the ORF1 and ORF2 start codons, in order to elucidate RNA vs non-RNA effects.

Thirdly, given the widespread use of retrotransposition reporters, we initially began our studies with a GFP-*LRE3* reporter construct. However, despite their widespread use, we note that a concern was raised about two decades ago regarding the possibility that these reporters may engage RNAi machinery due to the formation of dsRNAs stemming from the hybridization of RNAs produced from the sense-strand *L1* promoter and from the anti-sense strand CMV promoter that drives expression of the GFP cassette [42]. As such, it’s possible that some of the responses to retrotransposition reporters may be due to engagement of RNAi machinery and/or RNAi targeting of plasmid or endogenous *L1* RNA. Knockdown of *L1* expression through such a mechanism would also be consistent with downregulation of features like interferon signaling. However, it seems unlikely that this mechanism is the driver of the major pathway perturbations we observed, as (i) we detect an upregulation of plasmid and/or endogenous *L1* sequences following GFP-*LRE3* transfection and (ii) major perturbations were observed with other *L1* overexpression constructs lacking a reporter cassette (**Figure 7**). Nonetheless, future studies can further probe this question through the implementation of small RNA-sequencing to assess whether plasmid-derived siRNAs can be detected following transfection of reporters like GFP-*LRE3*. Finally, we note that for any given *L1* overexpression condition, we only assessed the impact on host cells at a single time point following transient transfection. As host responses are dynamic, it is possible that the relative differences in these responses, and interferon signaling in particular, may change over time and/or in response to constitutively sustained overexpression, as was done in the previously published RPE study. Ultimately, for all of these questions, this dataset collection may serve as a useful starting pointing for future studies aiming to further characterizing the impact of *L1* on host physiology.

## MATERIALS AND METHODS

### Re-processing of publicly available transcriptomic data

We leveraged publicly available mRNA-sequencing data for WI-38 human embryonic lung fibroblasts undergoing replicative senescence (BioProject PRJNA732700) [17] to characterize the effects of deep senescence on the repetitive element transcriptome. More specifically, we focused on transcriptomic data from fibroblasts that had undergone 45, 52, and 53 population doublings. Fastq files were first trimmed using fastp v0.20.1 [43] to (i) remove adapter sequences, (ii) hard trim the first 15 bases of each read to remove biased sequence composition, (iii) remove low quality bases, and (iv) remove reads shorter than 36 bases. Read quality for each sample was then inspected in the fastp quality report and separately using fastqc v0.11.9. Next, the GRCh38 primary human genome assembly and comprehensive gene annotation were obtained from GENCODE release 44 [44]. The trimmed reads were aligned to this reference genome using STAR v2.7.3a [45] with the following parameters: outFilterMultimapNmax 100, winAnchorMultimapNmax 100, and outFilterMismatchNoverLmax 0.04. Finally, the TEcount function in the TEtranscripts v2.1.4 [46] package was employed to obtain gene and repeat subfamily counts, using the GENCODE annotations to define gene boundaries and a repetitive element GTF file provided on the Hammell lab website (downloaded on September 14 2023 from https://labshare.cshl.edu/shares/mhammelllab/www-data/TEtranscripts/TE_GTF/GRCh38_GENCODE_rmsk_TE.gtf.gz) to define repeat element boundaries. We also leveraged publicly available mRNA-sequencing data for telomerase-immortalized retinal pigment epithelium-1 (RPE) cells harboring a doxycycline-inducible codon-optimized *L1* (*ORFeus*) or luciferase control (GSE119999) [32] to compare *L1* transcriptional signatures across cell types. This data was processed as described above, but using a modified reference genome and gene annotations containing the pDA093 *ORFeus* transgene as an additional contig.

Gene and repeat subfamily count files were loaded into R v4.3.3, which was used for all downstream analyses. To filter lowly expressed genes in each analysis, a counts-per-million (cpm) threshold corresponding to 10 reads in the median-length library was defined. Genes and repeat subfamilies were kept for analysis if they were expressed at levels surpassing this cpm threshold in at least as many samples as the smallest group. DESeq2 v1.42.1 [47] was used to identify significant (FDR < 0.05) differentially expressed genes and repeat subfamilies between groups or across continuous variables like population doublings. To visualize sample grouping patterns from the expression data, we carried out multidimensional scaling (MDS) analysis using a distance metric between samples based on Spearman’s rank correlation value (1-Rho), which was then provided to the ‘cmdscale’ R function.

### Plasmid Construction

The ks-99-GFP-LRE3 vector [18] was a gift from Dr. John Moran. pDA019 (Addgene plasmid # 131452; http://n2t.net/addgene:131452; RRID:Addgene_131452), pDA007 (Addgene plasmid # 131380; http://n2t.net/addgene:131380; RRID:Addgene_131380), pDA034 (Addgene plasmid # 131386; http://n2t.net/addgene:131386; RRID:Addgene_131386), and pDA077 (Addgene plasmid # 131388; http://n2t.net/addgene:131388; RRID:Addgene_131388) [32] were a gift from Dr. Kathleen Burns. The empty pcDNA3.1(+) backbone (Invitrogen cat. V79020) was a gift from the lab of Dr. Changhan David Lee at the University of Southern California Leonard Davis School of Gerontology. The empty pUC19 cloning vector was purchased from New England Biolabs (NEB; cat. N3041S).

We generated the ks-99-backbone plasmid, which lacks a retrotransposition cassette, to serve as an empty vector control for transfections with ks-99-GFP-LRE3. To do so, we digested ks-99-GFP-LRE3 with NotI and SalI, gel purified the ∼2900 bp fragment corresponding to the plasmid backbone using the Nucleospin Gel and PCR Cleanup Kit (Macherey-Nagel cat. 740609.50), blunted the fragment ends using the Quick Blunting Kit (NEB cat. E1201), and then self-ligated the fragment using the Quick Ligation Kit (NEB cat. M2200), generating ks-99-backbone. The overexpression constructs produced in this manuscript were generated by blunt-end cloning. To generate pcDNA3.1(+)_L1RP_ORF1-only, ORF1 was PCR-amplified from pDA077 with forward primer 5’-GCCGCCACCATGGGGAAAAAACAGAACAG-3’ and reverse primer 5’-GCAGTTTCTTCCTAGTCTTGATGG-3’ using AccuPrime *Pfx* DNA polymerase (Invitrogen cat. 12344024), the PCR product was purified with the Nucleospin Gel and PCR Cleanup Kit, and the fragment was ligated to EcoRV-digested, blunt-ended pcDNA3.1 using the Quick Ligation Kit. The pcDNA3.1(+)_L1RP_ORF2-only plasmid was generated with the same protocol but using forward primer 5’-GCCGCCACCATGACAGGATCAACTTCACAC-3’ and reverse primer 5’-TCCTGTGTCCATGTGATCTCATTG-3’ to amplify ORF2. The pcDNA3.1(+)_L1RP plasmid was similarly generated using forward primer 5’-CGCTCTAGCCCTGGAATGTG-3’ and reverse primer 5’-TGTCTGGATCTATTATACTCTAAG-3’ to amplify the full-length L1RP. The pUC19_L1RP plasmid was generated by ligating the same amplicon for pcDNA3.1(+)_L1RP to SmaI-digested, blunt-ended pUC19.

The plasmids generated in this study, ks-99-backbone (ID: 255270), pcDNA3.1(+)_L1RP_ORF1-only (ID: 255273), pcDNA3.1(+)_L1RP_ORF2-only (ID: 255274), pcDNA3.1(+)_L1RP (ID: 255272), and pUC19_L1RP (ID: 255271) are available through Addgene.

### Cell lines and cell culture conditions

Proliferative PDL9.74 IMR-90 (Coriell cat. I90-83, RRID: CVCL_0347) and PDL15 WI-38 (Coriell cat. AG06814-N, RRID: CVCL_0579) human embryonic fibroblasts were sourced from the Coriell Institute. Cells were maintained in Minimum Essential Medium (MEM) containing Earle’s salts (Corning cat. 15-010-CV), 15% fetal bovine serum (FBS, Sigma-Aldrich cat. F0926-500ML), 1X non-essential amino acids (NEAA, Quality Biological cat. 116-078-721), and 1X Penicillin-Streptomycin-Glutamine (Corning cat. 30-009-CI). Human Embryonic Kidney (HEK) 293T cells were purchased from ATCC (cat. CRL-3216) and were maintained in DMEM (Corning cat. 15-013-CV) containing 10% fetal bovine serum (FBS) and 1X Penicillin-Streptomycin-Glutamine. HeLa cells were a gift from the lab of Dr. Pinchas Cohen at the University of Southern California Leonard Davis School of Gerontology, and these were maintained in the same media as the HEK293T cells. Cells were cultured in a humidified incubator at 37°C and 5% CO_2_, subculturing cells once they reached ∼90% confluency. Cells were routinely tested for mycoplasma contamination using the PlasmoTest Mycoplasma Detection Kit (InvivoGen). The identities and purities of cells used in this study were verified using ATCC’s Human Cell STR Profiling Service (ATCC 135-XV).

### Retrotransposition assays

Retrotransposition assays following transient calcium phosphate transfection were carried out in HEK293T cells to confirm the mobilization-competency of the *LRE3* element on the ks-99-GFP-LRE3 plasmid. At 12-24 hours before transfection, 6-well plates were seeded with 250,000 cells per biological replicate and 6 replicates per group, and maintenance media was replaced 1-2 hours before transfection. We combined 3μg of DNA with calcium chloride (0.125 M final concentration) and HEPES-Buffered Saline (1X final concentration), using 0.1X Tris-EDTA buffer to normalize the plasmid volume. After allowing calcium phosphate crystals to form for ∼1 minute, the DNA-calcium phosphate mixes were added dropwise to each replicate, and plates were rocked back and forth to evenly distribute the mixes. After ∼24 hours, the cell culture media was replaced with fresh maintenance media. After ∼72 hours, cells were washed once with Dulbecco’s phosphate-buffered saline (DPBS, Corning cat. 21-031-CV), cells were detached using 0.05% trypsin (Corning cat. 25-052-CI), trypsin was neutralized using a two-fold higher volume of media, cells were pelleted (500xG, 5 minutes), and cells were resuspended in DPBS. The fraction of GFP^+^ cells in each sample was quantified on a MACSQuant Analyzer 10 (Miltenyi Biotec, 130-096-343), and flow cytometry data was analyzed in Flowlogic Solution 1.0. This experiment was repeated two times which yielded n = 12 independently transfected samples per group. Since the flow cytometer sensitivity was adjusted such that each empty vector control sample had ≤0.10% GFP^+^ cells, the proportions across experiments were directly combined, and statistical significance was reached if a Wilcoxon test yielded p < 0.05.

### Transfections and sample collections for ‘omics’ analyses

*E. coli* were cultured in LB Broth (Thermo Fisher Scientific) supplemented with 50 μg/mL carbenicillin to an optical density 600 (OD_600_) of 2 – 4. Plasmid extractions were carried out using the Nucleobond Xtra Midi Plus EF kit (Macherey-Nagel item no. 740422.50) following manufacturer recommendations. Plasmids were aliquoted and stored at −20°C until the time of transfection.

For paired transcriptomic and proteomic analyses, we washed IMR-90 fibroblasts twice with Dulbecco’s phosphate-buffered saline (DPBS, Corning cat. 21-031-CV), detached cells using 0.25% trypsin (HyClone cat. 95053-258), neutralized the trypsin using a two-fold higher volume of media, spun cells down (500xG, 5 minutes), resuspended cells in fresh media, and counted cells by trypan blue staining using a Countess II FL automated cell counter (Thermo Fisher) or a CellDrop Automated Cell Counter (DeNovix). The number of cells necessary for the experiment were then aliquoted, spun down, and washed once with DPBS. IMR-90 fibroblasts were transfected with ks-99-GFP-LRE3 or ks-99-backbone by electroporation using the Neon Transfection System (Invitrogen) with the following parameters: Buffer R for cell resuspension, 1500 V, 30 ms, and 1 pulse. Per reaction, we maintained a plasmid mass: cell number ratio of 5 μg: 2*10^6^ cells. We independently transfected 1.0 – 1.2 * 10^7^ fibroblasts per 100 mm tissue culture-treated petri dish (GenClone cat. 25-202), preparing 1 – 2 dishes to be used for each biological replicate downstream, with 6 biological replicates per experimental group. Immediately after transfection, cells were cultured in Penicillin-Streptomycin-free media and allowed to recover from the electroporation for ∼16 hours. Since cell viability following electroporation exhibited plate-to-plate variability, surviving cells were then detached by trypsinization, counted, and re-seeded at equal densities across biological replicates and experimental conditions. We subsequently defined this time point as “0 hours”.

IMR-90 fibroblasts were then prepared following previously described guidelines for generating conditioned media from senescent cells and quiescent control cells [19]. We implemented this protocol because (i) we hypothesized that *L1* overexpression might induce senescence-associated changes, based on the published literature, and (ii) this protocol utilizes serum deprivation to restrict proliferation and promote quiescence in order to eliminate differences due to proliferative versus non-proliferative states. More specifically, fibroblasts were re-seeded at a density of 6 * 10^6^ cells per 150 mm tissue culture-treated dish (Corning cat. 430599). Cells were cultured in 60 mL of 0.2% FBS media, which was replaced after 24 hours. At 48 hours, the media was removed, cells were washed twice with DPBS, and cells were fed with 60 mL of FBS-free and phenol red-free MEM media (Corning cat. 17-305-CV) containing Penicillin-Streptomycin-Glutamine and non-essential amino acids. At 72 hours, supernatants were collected in conicals, conicals were spun down (4000xG, 5 minutes, 4°C) to pellet cell debris, and supernatants were processed through Target2 0.45 μm polyvinylidene fluoride (PVDF) syringe filters (Thermo Scientific cat. F2500-5) to remove any remaining cellular debris. Filtered conditioned media was snap frozen in liquid nitrogen and stored at −80°C. Cells were washed with DPBS and detached with 0.05% trypsin (Corning cat. 25-052-CI), trypsin was neutralized with FBS-containing media, and cells were spun down. Cells were washed once with FBS-free and phenol red-free media, spun down, and resuspended again in that same media. Cell viability was assessed by trypan blue staining, and each sample was aliquoted into two tubes, saving 40% of the cells for proteomic analysis and 60% for transcriptomic analysis. Cells were spun down, the supernatants were discarded, and the cell pellets were snap frozen in liquid nitrogen and stored at −80°C. Cellular pellets for transcriptomic analysis were later lysed in TRIzol Reagent (Invitrogen) for downstream total RNA isolation (see below).

To more closely align the cell culture conditions to those used in published *L1* studies and ensure that major *L1*-induced pathway alterations were not solely a consequence of stress from the serum deprivation, we prepared additional transcriptomic samples for proliferating cells transfected with ks-99-GFP-LRE3 or ks-99-backbone cultured in rich, serum-containing maintenance media. IMR-90 fibroblasts were independently transfected as before and allowed to recover from the electroporation in Penicillin-Streptomycin-free maintenance media for 16-24 hours, plates were re-seeded at a density of 1 – 3 * 10^6^ cells per 100 mm dish containing standard maintenance media, and cells were collected in Trizol for downstream total RNA isolation 24 hours later with 4 biological replicates per experimental group. WI-38 fibroblasts were similarly prepared but with different electroporation parameters (1400 V, 20 ms, 2 pulses, and a plasmid mass: cell number ratio of 5 μg: 2*10^6^ cells), 4 biological replicates per experimental group, and collecting samples ∼72 hours after re-seeding. HEK293T cells were transfected through the calcium phosphate method described for the retrotransposition assays, with 3 biological replicates per experimental group and collecting samples ∼96 hours post-transfection. HeLa cells were also transfected through the described calcium phosphate approach, with 4 biological replicates per experimental group and collecting samples ∼48 hours post-transfection. N=3 HeLa samples were subsequently submitted for mRNA-sequencing.

To remove any potential impact of the retrotransposition reporter cassette and elucidate the impact of *L1* under various overexpression contexts, we prepared transcriptomic samples for proliferating IMR-90 cells transfected with pUC19_L1RP (expressing another active *L1* copy, *L1RP*, under its own internal promoter), pcDNA3.1(+)_L1RP_ORF1-only (expressing *L1RP* ORF1 under the CMV high-expression promoter), pcDNA3.1(+)_L1RP_ORF2-only (expressing *L1RP* ORF2 under the CMV promoter), pcDNA3.1(+)_L1RP (expressing full-length *L1RP* under the CMV promoter), pDA007 (expressing wild-type codon-optimized *ORFeus* under a doxycycline-inducible promoter), pDA034 (expressing codon-optimized *ORFeus*, with a mutation in the reverse transcriptase, under a doxycycline-inducible promoter), or empty vector controls. IMR-90 fibroblasts were prepared as described above under rich media conditions with 4 biological replicates per experimental group and collecting samples ∼72 hours after re-seeding. Importantly, 6 hours post-electroporation, Penicillin-Streptomycin-free media containing 2X doxycycline (Sigma cat. D9891-1G) was prepared and added to all plates in the *ORFeus* experiment for a final concentration of 1000 ng/mL. Standard rich media containing antibiotics, as well as 1000 ng/mL doxycycline, was used when re-seeding plates the following day and when replacing the media 48 hours post-reseeding. Finally, to ensure that our IMR-90 fibroblasts had the capacity to mount an interferon response, we also transfected them by electroporation with 5 μg of polyinosinic-polycytidylic acid (poly(I:C), Sigma cat. 528906-10MG) or water. After recovery from the electroporation, transfected IMR-90 fibroblasts were re-seeded on 6-well plates with 300,000 cells per well and 3 biological replicates per experimental group, and samples were collected ∼72 hours later.

All cells used were maintained below passage 30. Specifically, *LRE3* multi-omics samples on quiescent IMR-90 cells were prepared from cells at passage 11, *LRE3* samples in proliferating IMR-90 cells were prepared at passage 11, *LRE3* samples in proliferating WI-38 cells were prepared at passage 20, *LRE3* samples in proliferating HEK293T cells were prepared at passage 14, *LRE3* samples in proliferating HeLa cells were prepared at passage 18, *L1RP* (native promoter) samples in proliferating IMR-90 cells were prepared at passage 8, *L1RP* (CMV promoter) samples in proliferating IMR-90 cells were prepared at passage 8, *ORFeus* samples in proliferating IMR-90 cells were prepared at passage 8, and poly(I:C) samples in proliferating IMR-90 cells were prepared at passage 15.

### RNA extractions, PCR and RT-qPCR validations, and mRNA sequencing

RNA was extracted using the Direct-zol RNA Miniprep kit (Zymo Research cat. R2052) following manufacturer recommendations. To confirm *LRE3* overexpression from ks-99-GFP-LRE3, complementary DNA (cDNA) was first generated for each RNA sample using the Maxima H Minus cDNA Synthesis Master Mix with dsDNase (Thermo Scientific cat. M1682), following manufacturer instructions and including reverse transcriptase negative (RT-) controls. Plasmid-specific sense-strand *LRE3* overexpression was verified by endpoint PCR using MyTaq HS Red Mix (Meridian Bioscience cat. BIO-25048) with forward/reverse primers JB8046/JB9501 (5’-ACCCAACACCCGTGCGTTTTATT-3’ / 5’-TGGAGTACAACTACAACAGCCACAACGTCT-3’), which can produce 1401 bp and 499 bp amplicons, or with primers eGFP-spliced-2R/ks99-pcr2R (5’-CTTGTACAGCTCGTCCATGCC-3’ / 5’-AATGGGCGTGGATAGCGG-3’), which can produce 1871 bp and 969 bp amplicons. These primer pairs flank the intron in the GFP reporter cassette and produce a larger and a smaller amplicon depending on whether the intron is unspliced or spliced. Importantly, intron splicing should only occur when transcription occurs on the sense strand; thus, the presence of the smaller amplicon corresponds to transcription from the *LRE3* internal promoter on the sense strand and not transcription from the CMV promoter that drives GFP expression on the anti-sense strand. For HEK293T samples, *LRE3* overexpression was assessed by visually inspecting all samples under a microscope and checking for GFP^+^ cells, which would indicate successful transcription and mobilization of the *LRE3* copy on the ks-99-GFP-LRE3 plasmid.

*L*1 overexpression in the remaining samples was assessed by reverse transcription followed by quantitative polymerase chain reaction (qPCR). Three-step qPCR was carried out using the SensiFAST SYBR No-ROX Kit (Thomas Scientific cat. BIO-98020) in a magnetic induction cycler (MIC) (Bio Molecular Systems cat. BMS-MIC-2) with these steps: polymerase activation (95°C, 2 minutes) and 40 cycles of denaturation (95°C, 5 seconds), annealing (60°C, 10 seconds), and extension (72°C, 20 seconds). For pUC19_L1RP, since we could not verify plasmid-specific *L1* overexpression, we quantified relative expression of evolutionarily-young *L1* elements, including *L1RP*, using previously published primers targeting the 5’UTR (5’-GCCAAGATGGCCGAATAGGA-3’ / 5’-AAATCACCCGTCTTCTGCGT-3’), ORF1 (5’-ACCTGAAAGTGACGGGGAGA-3’ / 5’-CCTGCCTTGCTAGATTGGGG-3’), and ORF2 (5’-CAAACACCGCATATTCTCACTCA-3’ / 5’-CTTCCTGTGTCCATGTGATCTCA-3’) [9]. For pcDNA3.1(+)_L1RP, pcDNA3.1(+)_L1RP_ORF1-only, and pcDNA3.1(+)_L1RP_ORF2-only, plasmid-specific overexpression was assessed using primers targeting the CMV promoter and the *L1RP* 5’UTR (5’-CGCTCTAGCCCTGGAATGTG-3’ / 5’-AAATCACCGTCTTCTGCGT-3’), ORF1 (5’-AAACTTAAGCTTGGTACCGAGC-3’ / 5’-GCGTTCCTTTGGAGGAGGAG-3’), or ORF2 (5’-CTAGTCCAGTGTGGTGGAATTC-3’ / 5’-ATCCAACTTGCCAGTCTGTGTC-3’). For pDA007 and pDA034, *ORFeus* overexpression was assessed using primers targeting *ORFeus* ORF1 (5’-AGAACGACTTCGACGAGCTG-3’ / 5’-CTTCAGGCACTTCTCGGTGT-3’) and *ORFeus* ORF2 (5’-CAGAACCGCGATATCGACCA-3’ / 5’-AGTTCTCCCAGCACCACTTG-3’). To control for technical differences in loading volumes, we also measured the relative expression of reference genes *UBC* (5’-ACTCTGCACTTGGTCCTGC-3’ / 5’-CAACTTTATTGAAAGGAAAGTGCAA-3’), *ABL1* (5’-GCCTCCTTCTTCCACTTCTC-3’ / 5’-ATGCCCTTCCCGAAATGC-3’), and *HSP90AB1* (5’-AAGAGAGCAAGGCAAAGTTTGAG-3’ / 5’-TGGTCACAATGCAGCAAGGT-3’). Changes in gene expression were quantified using the delta-delta Ct method, using the geometric mean of the Ct values for all three reference genes as a loading control, and normalizing expression values to the median of the control group in each experiment. Statistical analysis was carried out in R using the Wilcox.test function.

The integrity of RNA samples were evaluated using an Agilent High Sensitivity RNA ScreenTape assay (Agilent Technologies) or by Agilent 2100 Bioanalyzer at Novogene (Sacramento, California). Of the 6 biological replicates per group in the paired transcriptome/proteome experiment, 4-5 samples with eRIN scores ranging between 8.8 and 9.4 were selected for sequencing. Across all of the follow-up experiments in standard, serum-containing media, all tested samples had eRIN scores ranging between 8.2 and 9.6. Because of Tapestation System maintenance, the integrities of HeLa samples were evaluated by Novogene and passed quality control. Novogene carried out unstranded mRNA library preparation and sequencing on the NovaSeq 6000 or NovaSeq X Plus platform as paired-end 150 bp reads. The raw FASTQ reads have been deposited to SRA under BioProject PRJNA1174135.

### Protein digestion and desalting

Frozen cellular pellets and frozen conditioned media were shipped to the USC-Buck Institute Nathan Shock Center Cellular Senescence and Beyond Core (CSBC) for proteomic analysis. Conditioned media (30 mL per sample) from either empty vector (N=4) or *LRE3* overexpression (N=4) IMR-90 cultures were concentrated using 3 kDa molecular cut-off filters (Millipore Sigma, Burlington, MA) and protein concentrations were determined using

Bicinchoninic Acid (BCA) assay (Thermo Fisher Scientific, Waltham, MA). Cell pellets from either empty vector (N=6) or *LRE3* overexpression (N=6) IMR-90 cultures were solubilized in 100 µL of 0.5% sodium dodecyl sulfate (SDS) in 100 mM Triethylamonium bicarbonate (TEAB) with 1X protease inhibitor cocktail (PIC). Protein concentrations were determined using BCA assay.

Samples originating from cell pellets (∼100 µg) were brought to the same overall volume of 100 µL with water, while secretome samples (∼100 µg) were prepared directly. All samples were reduced using 20 mM dithiothreitol in 50 mM TEAB at 50°C for 10 minutes, cooled to room temperature (RT) and held at RT for 10 minutes, and alkylated using 40 mM iodoacetamide in 50 mM TEAB at RT in the dark for 30 minutes. Samples were acidified with 12% phosphoric acid to obtain a final concentration of 1.2% phosphoric acid. S-Trap buffer consisting of 90% methanol in 100 mM TEAB at pH ∼7.1 was added and samples were loaded onto S-Trap micro spin columns (Protifi). The entire sample volume was spun through the S-Trap micro spin columns at 4,000 x g and RT, binding the proteins to the micro spin columns. Subsequently, S-Trap micro spin columns were washed twice with S-Trap buffer at 4,000 x g and RT and placed into clean elution tubes. Samples were incubated for one-hour at 47°C with sequencing grade trypsin (Promega, San Luis Obispo, CA) dissolved in 50 mM TEAB at a 1:25 (w/w) enzyme:protein ratio. An additional aliquot of trypsin dissolved in 50 mM TEAB was added and samples were digested overnight at 37°C.

Peptides were sequentially eluted from micro S-Trap spin columns with 50 mM TEAB, 0.5% formic acid (FA) in water, and 50% acetonitrile (ACN) in 0.5% FA. After centrifugal evaporation, samples were resuspended in 0.2% FA in water and desalted with Oasis 10-mg Sorbent Cartridges (Waters, Milford, MA). The desalted elutions were then subjected to an additional round of centrifugal evaporation and re-suspended in 0.1% FA in water at a final concentration of 1 µg/µL. Eight microliters of each sample were diluted with 2% ACN in 0.1% FA to obtain a concentration of 400 ng/µL. One microliter of indexed Retention Time Standard (iRT, Biognosys, Schlieren, Switzerland) was added to each sample, thus bringing up the total volume to 20 µL [48].

### Mass spectrometric analysis (secretome)

Reverse-phase high-performance liquid chromatography (HPLC)-MS/MS analyses were performed on a Dionex UltiMate 3000 system coupled online to an Orbitrap Exploris 480 mass spectrometer (Thermo Fisher Scientific, Bremen, Germany). The solvent system consisted of 2% ACN, 0.1% FA in water (solvent A) and 80% ACN, 0.1% FA in ACN (solvent B). Digested peptides (800 ng) were loaded onto an Acclaim PepMap 100 C_18_ trap column (0.1 x 20 mm, 5 µm particle size; Thermo Fisher Scientific) over 5 minutes at 5 µL/min with 100% solvent A. Peptides were eluted on an Acclaim PepMap 100 C_18_ analytical column (75 µm x 50 cm, 3 µm particle size; Thermo Fisher Scientific) at 300 nL/min using the following gradient: linear from 2.5% to 24.5% of solvent B in 125 min, linear from 24.5% to 39.2% of solvent B in 40 min, up to 98% of solvent B in 1 min, and back to 2.5% of solvent B in 1 min. The column was re-equilibrated for 30 min with 2.5% of solvent B, and the total gradient length was 210 min. Each sample was acquired in data-independent acquisition (DIA) mode [49–51]. Full MS spectra were collected at 120,000 resolution (Automatic Gain Control (AGC) target: 3e6 ions, maximum injection time: 60 ms, 350-1,650 *m/z*), and MS2 spectra at 30,000 resolution (AGC target: 3e6 ions, maximum injection time: Auto, Normalized Collision Energy (NCE): 30, fixed first mass 200 *m/z*). The isolation scheme consisted of 26 variable windows covering the 350-1,650 *m/z* range with an overlap of 1 *m/z* (see **Supplementary Table S1D**) [50].

### Mass spectrometric analysis (intracellular protein)

LC-MS/MS analyses were performed on a Dionex UltiMate 3000 system coupled to an Orbitrap Eclipse Tribrid mass spectrometer (both from Thermo Fisher Scientific, San Jose, CA). The solvent system consisted of 2% ACN, 0.1% FA in water (solvent A) and 98% ACN, 0.1% FA in water (solvent B). Proteolytic peptides (600 ng) were loaded onto an Acclaim PepMap 100 C_18_ trap column (0.1 x 20 mm, 5-µm particle size; Thermo Fisher Scientific) for 5 min at 5 µL/min with 100% solvent A. Peptides were eluted on an Acclaim PepMap 100 C_18_ analytical column (75 µm x 50 cm, 3 µm particle size; Thermo Fisher Scientific) at 300 nL/min using the following gradient of solvent B: 2% for 5 min, linear from 2% to 20% in 125 min, linear from 20% to 32% in 40 min, up to 80% in 1 min, 80% for 9 min, and down to 2% in 1 min. The column was equilibrated with 2% of solvent B for 29 min, with a total gradient length of 210 min. All samples were acquired in DIA mode. Full MS spectra were collected at 120,000 resolution (AGC target: 3e6 ions, maximum injection time: 60 ms, 350-1,650 m/z), and MS2 spectra at 30,000 resolution (AGC target: 3e6 ions, maximum injection time: Auto, NCE: 27, fixed first mass 200 m/z). The isolation scheme consisted of 26 variable windows covering the 350-1,650 m/z range with an overlap of 1 *m/z* (see **Supplementary Table S1D**) [50].

### DIA-MS Data Processing and Statistical Analysis

DIA data was processed in Spectronaut (versions 15.7.220308.50606 and 16.2.220903.53000) using directDIA. Data extraction parameters were set as dynamic and non-linear iRT calibration with precision iRT was selected. Data was searched against the *Homo Sapiens* reference proteome with 20,380 entries (UniProtKB-SwissProt), accessed on 01/29/2021. Trypsin/P was set as the digestion enzyme and two missed cleavages were allowed. Cysteine carbamidomethylation was set as a fixed modification while methionine oxidation and protein N-terminus acetylation were set as dynamic modifications. Identification was performed using 1% precursor and protein q-value. Quantification was based on the peak areas of extracted ion chromatograms (XICs) of 3 – 6 MS2 fragment ions, specifically b- and y-ions, with q-value sparse data filtering and iRT profiling applied (**Supplementary Table S1H**). Local normalization was applied for intracellular protein analysis (i.e., cell pellets), but not for secretome analysis. We note that 1 of the empty vector secretome samples was excluded from the downstream analyses, as it was objectively assessed as an outlier (different profile from the remaining control samples as was determined by partial least squares discriminant analysis). Consequently, only N=3 instead of N=4 secretome samples were analyzed for the control group for robustness. Differential protein expression analysis comparing 1) *LRE3* overexpression to empty vector was performed using a paired t-test, and p-values were corrected for multiple testing, using the Storey method [52]. Specifically, group wise testing corrections were applied to obtain q-values. For intracellular protein analysis, protein groups with at least two unique peptides, q-value < 0.05, and absolute log_2_(fold-change) > 0.2 were called as significantly-altered (**Supplementary Table S1F**). For secretome analysis, protein groups with at least two unique peptides, q-value < 0.05, and absolute log_2_(fold-change) > 0.58 were called as significantly-altered (**Supplementary Table S1H**).

### RNA-seq read trimming, mapping, and quantification

Fastq files for all of the samples generated in this study were trimmed, mapped, and quantified as described above for the re-analysis of publicly available data, except that a slightly modified reference genome and gene annotations containing the *L1* overexpression plasmid were used. For the *LRE3* datasets, the ks-99-GFP-LRE3 plasmid sequence was appended to the GRCh38 primary human genome assembly from GENCODE. For the analyses with *L1RP* or its individual ORFs under the CMV promoter, the pcDNA3.1(+)_L1RP, pcDNA3.1(+)_L1RP_ORF1-only, and pcDNA3.1(+)_L1RP_ORF2-only plasmid sequences were all appended to the GRCh38 assembly. For the analyses with *L1RP* under its own internal promoter, the pUC19_L1RP plasmid sequence was appended to the GRCh38 assembly. For the analysis with *ORFeus* and its mutant, the pDA007 plasmid sequence was appended to the GRCh38 assembly. The *L1* 5’UTR and ORF coordinates were also included in the corresponding gene GTF annotations from GENCODE release 44. We did this to assess plasmid-specific *L1* overexpression by RNA-seq, as a complementary analysis to our PCR-based or fluorescence-based overexpression validation approaches. We note, however, that precise quantification of evolutionarily young *L1* copies (like *LRE3* and *L1RP*) is extremely difficult [53], and reads originating from plasmid *L1* copies may be re-distributed to the global *L1Hs* pool by the TEtranscripts v2.1.4 [46] algorithm, or vice versa, and TE reads from any copy may be omitted from analysis if they map to too many loci. Biologically, the potential generation of truncated transcripts from a single *L1* copy due to the presence of internal canonical and noncanonical polyadenylation signals also complicates the biological and computational study of *L1* [33]. As such, these quantifications should be interpreted in conjunction with the complementary validation analyses. For the poly(I:C) analysis, the unmodified reference genome and annotations were used.

Gene and repeat subfamily count files were loaded into R v4.3.3, which was used for all downstream analyses. To filter lowly expressed genes in each analysis, a counts-per-million (cpm) threshold corresponding to 10 reads in the median-length library was defined. Genes and repeat subfamilies were kept for analysis if they were expressed at levels surpassing this cpm threshold in at least as many samples as the smallest group. For proliferating IMR-90 fibroblasts overexpressing *LRE3*, surrogate variables were estimated with the ‘svaseq’ function [54] in the R package ‘sva’ v3.50.0, and they were regressed out using the ‘removeBatchEffect’ function in the R package limma v3.58.1 [55]. DESeq2 v1.42.1 [47] was used to identify significant (FDR < 0.05) differentially expressed genes and repeat subfamilies between groups. To visualize sample grouping patterns from the expression data, we carried out multidimensional scaling (MDS) analysis using a distance metric between samples based on Spearman’s rank correlation value (1-Rho), which was then provided to the ‘cmdscale’ R function.

### Functional enrichment analyses

We used the Gene Set Enrichment Analysis (GSEA) paradigm as implemented in the R package clusterProfiler v4.10.1 [56]. Hallmark gene sets [22] were obtained from the Molecular Signature Database release 2024.1.Hs [20]. Reactome v92 pathway gene sets were obtained directly from the Reactome website [21]. For a subset of analyses, GSEA was carried out only on Hallmark and Reactome interferon-related gene sets. We also obtained several aging- and senescence-associated gene sets, including core senescence-associated secretory phenotype (SASP) factors [27], SenMayo [26], CellAge build 3 [25], aging-related genes derived from the Genotype-Tissue Expression project [24], differentially expressed genes/repeats identified in our recent re-analysis [16] of human primary aging fibroblasts [57], and differentially expressed genes/repeats identified in our analysis of WI-38 fibroblasts undergoing deep senescence [17]. We also highlighted senescence and interferon-related genes identified by De Cecco *et al.* [9].

For gene set enrichment analysis of transcriptomic changes, the DESeq2 v1.42.1 Wald-statistic was used to generate a ranked list of gene symbols and repeats. For gene set enrichment analysis of proteomic and secretomic changes, the product of the log_2_(fold change) and -log_10_(Q-value) was used to generate a ranked list of proteins, which were then mapped to their corresponding gene symbol. All gene sets with an FDR < 0.05 were considered significant. For plots with a single analysis, the top 5 downregulated and top 5 upregulated gene sets were plotted, at most.

### Multi-contrast pathway enrichment analyses

Multi-contrast pathway enrichment was carried out across “-omic” layers or across *L1* construct-specific transcriptomic profiles using the mitch v1.14.0 R package [40]. We used a subset of the gene sets used for GSEA, including Hallmark gene sets [22] obtained from the Molecular Signature Database release 2024.1.Hs [20] and Reactome v92 pathway gene sets obtained directly from the Reactome website [21]. For multi-contrast pathway enrichment analysis of transcriptomic changes, the DESeq2 v1.42.1 Wald-statistic was used to generate a ranked list of gene symbols and repeats. For multi-contrast pathway enrichment analysis of proteomic and secretomic changes, the product of the log_2_(fold change) and -log_10_(Q-value) was used to generate a ranked list of proteins, which were then mapped to their corresponding gene symbol. Gene sets with an adjusted MANOVA (Multivariate Analysis of Variance) p-value < 0.05 were considered significant.

### Cell cycle stage profiling using propidium iodide staining

Cell cycle statuses of *LRE3* or empty vector transfected cells were determined by propidium iodide (PI) staining. After the electroporation recovery period, cells were detached, and ∼500,000 – 600,000 cells per well were seeded on 6-well plates containing maintenance media, with 3 biological replicates per group. After 24 hours, cells were detached, spun down (300xG, 5 minutes), washed once in DPBS, fixed by drop-wise adding pre-chilled 70% ethanol, and fixed cells were stored at −20°C for a minimum of 12 hours. Then, cells were spun down (500xG, 5 minutes, 4°C), cells were washed once with ice-cold DPBS, resuspended in labeling buffer, and stained for 30 minutes in the dark. Staining buffer was prepared in DPBS with the following final concentrations: 50 μg/mL propidium iodide (Alfa Aesar cat. J66584), 100 μg/mL

PureLink RNase A (Invitrogen cat. 12091021), and 0.05% Triton X-100 (MP cat. 194854). Flow cytometry analysis was carried out on a MACSQuant Analyzer 10 (Miltenyi Biotec, 130-096-343) and flow cytometry data was analyzed in Flowlogic Solution 1.0. This experiment was independently repeated in two batches, yielding n = 6 independently transfected samples per group. Each set of cell cycle phase proportions were normalized to the median of the control group in each experiment, the values across experiments were combined, and statistical significance was reached if a Wilcoxon test yielded p < 0.05.

### Other software versions

Analyses were conducted using R version 4.3.3 and code was re-run independently to check for reproducibility.

## Data and code availability

RNA-seq data have been deposited at the Sequence Read Archive (SRA) under BioProject PRJNA1174135. Raw data and complete mass spectrometry data sets have been uploaded to the Mass Spectrometry Interactive Virtual Environment (MassIVE) repository, developed by the Center for Computational Mass Spectrometry at the University of California San Diego, and can be downloaded using the following link: https://massive.ucsd.edu/ProteoSAFe/private-dataset.jsp?task=6688d0c9925749d187e256d24c4fe30c (MassIVE ID number: MSV000103003; ProteomeXchange ID: PXD083088). Original DNA gel images, RT-qPCR Ct data, and raw flow cytometry data have been deposited at Figshare at [https://doi.org/10.6084/m9.figshare.33202578]. All scripts used to analyze the data in this manuscript are available on the Benayoun Lab GitHub at https://github.com/BenayounLaboratory/LINE1_overexpression.

## ACKNOWLEDGMENTS

We are saddened by the passing of Dr. Judith Campisi and gratefully acknowledge her valuable contribution to this work before her passing. Some illustrations used to assemble this manuscript’s figures were obtained from the NIAID NIH BIOART Source (bioart.niaid.nih.gov) or Bioicons (bioicons.com). We thank Dr. Hassy Cohen for the generous gift of HeLa cells used in our study, and we thank Dr. John Moran, Dr. Kathleen Burns, and Dr. Changhan David Lee for the generous gift of plasmids that were also used in this study. We acknowledge Addgene for facilitating the transfer of plasmids, including the ones generated as part of this manuscript.

## SOURCES OF FUNDING

This work was supported by fellowship DGE-1842487 from the National Science Foundation Graduate Research Fellowship Program, predoctoral fellowship of T32 AG052374 from NIH/NIA, and a University of Southern California Provost Fellowship to J.I.B.; a Biology of Aging PhD fellowship from the USC Leonard Davis School of Gerontology to E.T.; and by NIGMS R35 GM142395 award to B.A.B. This work was also made possible by a pilot award under NIA grant P30 AG068345 (USC-Buck Institute Nathan Shock Center of Excellence) and by the NIH-OD instrumentation grant S10 OD028654 (to B.S.) for the Orbitrap Eclipse Tribrid mass spectrometry system. The funders had no role in study design, data collection and analysis, decision to publish, or preparation of the manuscript.

## ETHICS AND CONSENT TO PARTICIPATE DECLARATIONS

Not applicable.

## AUTHOR CONTRIBUTIONS

Juan I. Bravo: Conceptualization, Data curation, Investigation, Formal analysis, Visualization, Writing – original draft, Writing – review & editing. Eyael Tewelde: Investigation, Formal analysis, Validation, Writing – original draft, Writing – review & editing. Christina D. King: Investigation, Formal analysis, Writing – original draft, Writing – review & editing. Nayana S. Tellakula: Validation. Birgit Schilling: Conceptualization, Formal analysis, Writing – original draft, Writing – review & editing, Supervision, Funding acquisition. Bérénice A. Benayoun: Conceptualization, Data curation, Formal analysis, Visualization, Writing – original draft, Writing – review & editing, Supervision, Funding acquisition.

## DECLARATION OF INTERESTS

The authors declare that they have no competing interests. The content is solely the responsibility of the authors and does not necessarily represent the official views of the National Institutes of Health.

## SUPPLEMENTARY FIGURE TITLES AND LEGENDS

**Supplementary Fig. S1. *LRE3* is a retrotransposition-competent *L1* copy (A)** A diagram illustrating how the retrotransposition-competence of an *L1* (*LRE3*) copy was assessed in HEK293T cells. **(B)** The flow cytometry gating strategy for the *L1* transposition assay. **(C)** Quantification of the GFP+ fraction of HEK293T cells (N=12 per group).

**Supplementary Fig. S2. Quality control of the samples used for multi-omic profiling. (A)** Plasmid-specific *LRE3* overexpression was assessed by endpoint RT-PCR of empty vector and *LRE3* overexpressing quiescent IMR-90 fibroblasts. PCR reactions were carried out with (RT+) or without (RT-) reverse transcription in N = 6 samples per group, and N = 4-6 of these were used for multi-omic profiling. We note that 1 of the empty vector secretome samples was excluded from all downstream analyses, as it was objectively assessed as an outlier (different profile from the remaining control samples as was determined by partial least squares discriminant analysis). Consequently, only N=3 instead of N=4 secretome samples for the control group were utilized for robustness (see **methods**). A table showing samples utilized for each-omics analysis is also shown. **(B)** Heatmap of RNA-seq quantification of *LRE3* features in transfected cells. **(C)** Multidimensional scaling (MDS) analysis of the repetitive element transcriptome across samples.

Supplementary Fig. S3. *L1*-induced changes show opposite directionality with changes seen in organismal aging and cellular senescence. (A) A diagram illustrating how *LRE3*-induced multi-omic changes were compared to known senescence- and aging-induced features. GSEA analysis with gene sets for senescence- and aging-induced features in **(B)** the transcriptome, cellular proteome, and secretome of *LRE3* overexpressing fibroblasts and **(D)** the transcriptomes of fibroblasts in deep replicative senescence (GSE175533; Figure 1). Gene sets with FDR < 0.05 were considered significant. **(C)** Expression of deep senescence-associated genes identified by De Cecco *et al.*, 2019 in *LRE3* overexpressing fibroblasts. SASP: Senescence-Associated Secretory Phenotype, FDR: False Discovery Rate, NES: Normalized Enrichment Score.

**Supplementary Fig. S4. (A)** The flow cytometry gating strategies for the propidium iodide-based cell cycle assay.

**Supplementary Fig. S5. Transient *L1* overexpression leads to dampening of interferon pathways.** GSEA for (**A**) Reactome interferon signaling gene set regulation and (**B)** Hallmark interferon gamma gene set regulation in *LRE3* overexpressing fibroblasts across ‘omic’ layers. GSEA: Gene Set Enrichment Analysis, NES: Normalized Enrichment Score.

**Supplementary Fig. S6. Quality control of proliferating IMR-90 fibroblasts overexpressing *LRE3* used for mRNA-sequencing. (A)** Plasmid-specific *LRE3* overexpression was assessed by endpoint RT-PCR of empty vector and *LRE3*-overexpressing, proliferating IMR-90 fibroblasts. PCR reactions were carried out with (RT+) or without (RT-) reverse transcription in N = 4 samples per group. Those same samples were submitted for sequencing. **(B)** RNA-seq quantification of *LRE3* features in transfected cells. **(C)** Multidimensional scaling (MDS) analysis of the gene and repetitive element transcriptomes across samples.

**Supplementary Fig. S7. Quality control of proliferating WI-38 fibroblasts overexpressing *LRE3* used for mRNA-sequencing. (A)** Plasmid-specific *LRE3* overexpression was assessed by endpoint RT-PCR of empty vector and *LRE3* overexpressing proliferating WI-38 fibroblasts. PCR reactions were carried out with (RT+) or without (RT-) reverse transcription in N = 4 samples per group. Those same samples were submitted for sequencing. **(B)** RNA-seq quantification of *LRE3* features in transfected cells. **(C)** Multidimensional scaling (MDS) analysis of the gene and repetitive element transcriptomes across samples.

**Supplementary Fig. S8. Quality control of proliferating IMR-90 fibroblasts overexpressing *L1RP* used for mRNA-sequencing. (A)** A diagram illustrating how the transcriptomic impact of *L1RP* overexpression in proliferating IMR-90 fibroblasts were assessed. **(B)** RT-qPCR quantification of global, evolutionarily-young *L1* 5’UTR, ORF1 and ORF2 levels in empty vector and *L1RP* transfected fibroblasts. N=4 samples per group were tested, and those same samples were submitted for sequencing. **(C)** RNA-seq quantifications of endogenous, evolutionarily-young *L1Hs* levels. **(D)** Multidimensional scaling (MDS) analysis of the gene and repetitive element transcriptomes across samples.

**Supplementary Fig. S9. Quality control of proliferating IMR-90 fibroblasts overexpressing CMV-*L1RP* used for mRNA-sequencing. (A)** A diagram illustrating how the transcriptomic impacts of *CMV*-driven full-length *L1*, ORF1 and ORF2 overexpression in proliferating IMR-90 fibroblasts were assessed. **(B)** RT-qPCR quantification of plasmid-specific full-length *L1*, ORF1 and ORF2 overexpression in empty vector and *L1RP* transfected fibroblasts. N=4 samples per group were tested, and those samples were submitted for sequencing. **(C)** RNA-seq quantifications of plasmid *L1RP* features. **(D)** Multidimensional scaling (MDS) analysis of the gene and repetitive element transcriptomes across samples.

**Supplementary Fig. S10. Quality control of proliferating IMR-90 fibroblasts overexpressing *ORFeus* used for mRNA-sequencing. (A)** A diagram illustrating how the transcriptomic impact of codon-optimized *L1* (*ORFeus*), with or without reverse transcriptase, in proliferating IMR-90 fibroblasts was assessed. **(B)** RT-qPCR quantification of codon-optimized ORF1 and ORF2 expression in empty vector and *ORFeus*-transfected fibroblasts. N=4 samples per group were tested, and those samples were submitted for sequencing. **(C)** RNA-seq quantifications of plasmid-specific *ORFeus* features. **(D)** Multidimensional scaling (MDS) analysis of the gene and repetitive element transcriptomes across samples.

**Supplementary Fig. S11. Quality control of proliferating HeLa cells overexpressing *LRE3* used for mRNA-sequencing. (A)** Plasmid-specific *LRE3* overexpression was assessed by endpoint RT-PCR of empty vector and *LRE3*-overexpressing proliferating HeLa cells. PCR reactions were carried out with (RT+) or without (RT-) reverse transcription in N = 4 samples per group, and N=3 samples per group were submitted for sequencing. **(B)** RNA-seq quantification of *LRE3* features in transfected cells. **(C)** Multidimensional scaling (MDS) analysis of the gene and repetitive element transcriptomes across samples.

**Supplementary Fig. S12. Quality control of proliferating HEK293T cells overexpressing *LRE3* used for mRNA-sequencing. (A)** RNA-seq quantification of *LRE3* features in transfected cells. **(B)** Multidimensional scaling (MDS) analysis of the gene and repetitive element transcriptomes across samples.

**Supplementary Fig. 13. Quality control of RPE *ORFeus* and IMR-90 poly(I:C) transcriptomic datasets. (A)** Multidimensional scaling (MDS) analysis of the gene and repetitive element transcriptomes of RPE cells overexpressing *ORFeus* (GSE119999). **(B)** A diagram illustrating how the transcriptomic impact of the viral mimetic poly(I:C) was assessed in IMR-90 fibroblasts. **(C)** Multidimensional scaling (MDS) analysis of the gene and repetitive element transcriptomes across samples.

**Supplementary Fig. S14. (A)** Differential expression of MSigDB Hallmark interferon alpha and interferon gamma pathway genes across the various *L1* overexpression and poly(I:C) exposure datasets.

## Notes

### Competing Interest Statement

The authors have declared no competing interest.

