## Supplementary figures for "Acute *LINE-1* overexpression rewires the transcriptome, proteome and secretome of primary human fibroblasts"

Figure S1

**A** Scheme for assessing the functionality of a *LINE-1* (*LRE3*) transposition reporter in HEK293T cells

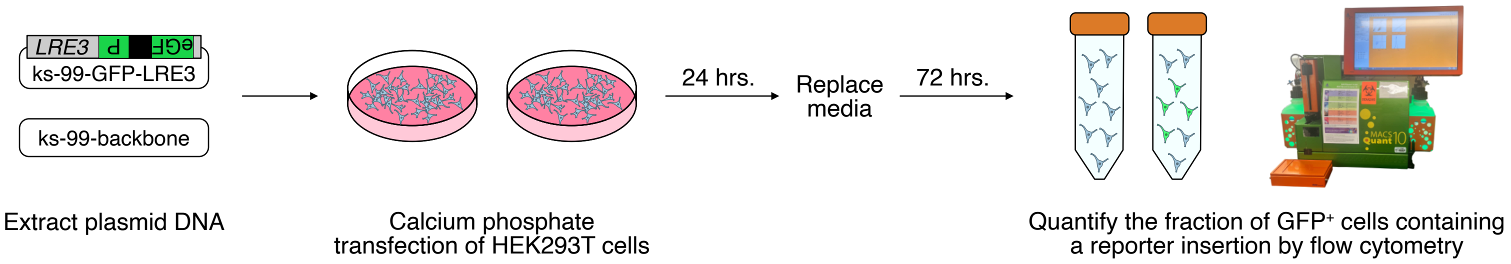


**B** Flow cytometry gating strategy for the transposition assay (*LRE3* example)

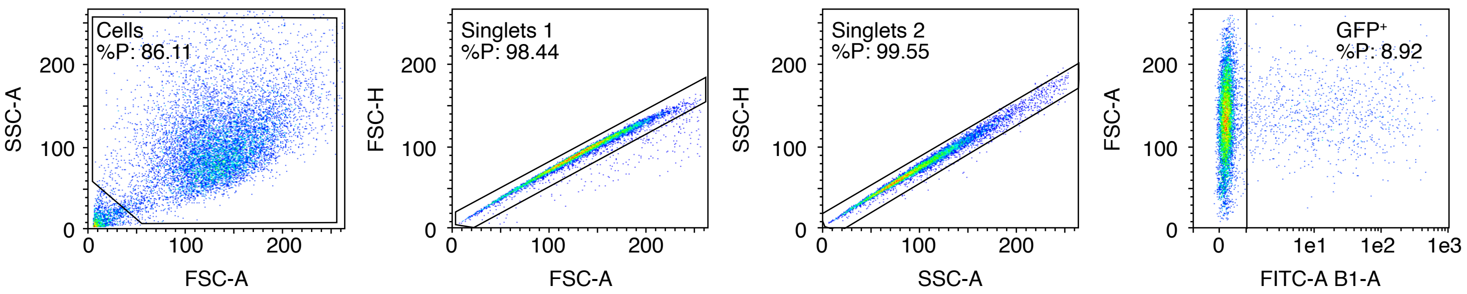


**C** Quantification of the GFP⁺ fraction

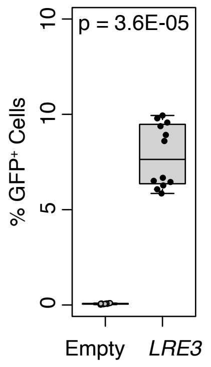

Figure S2

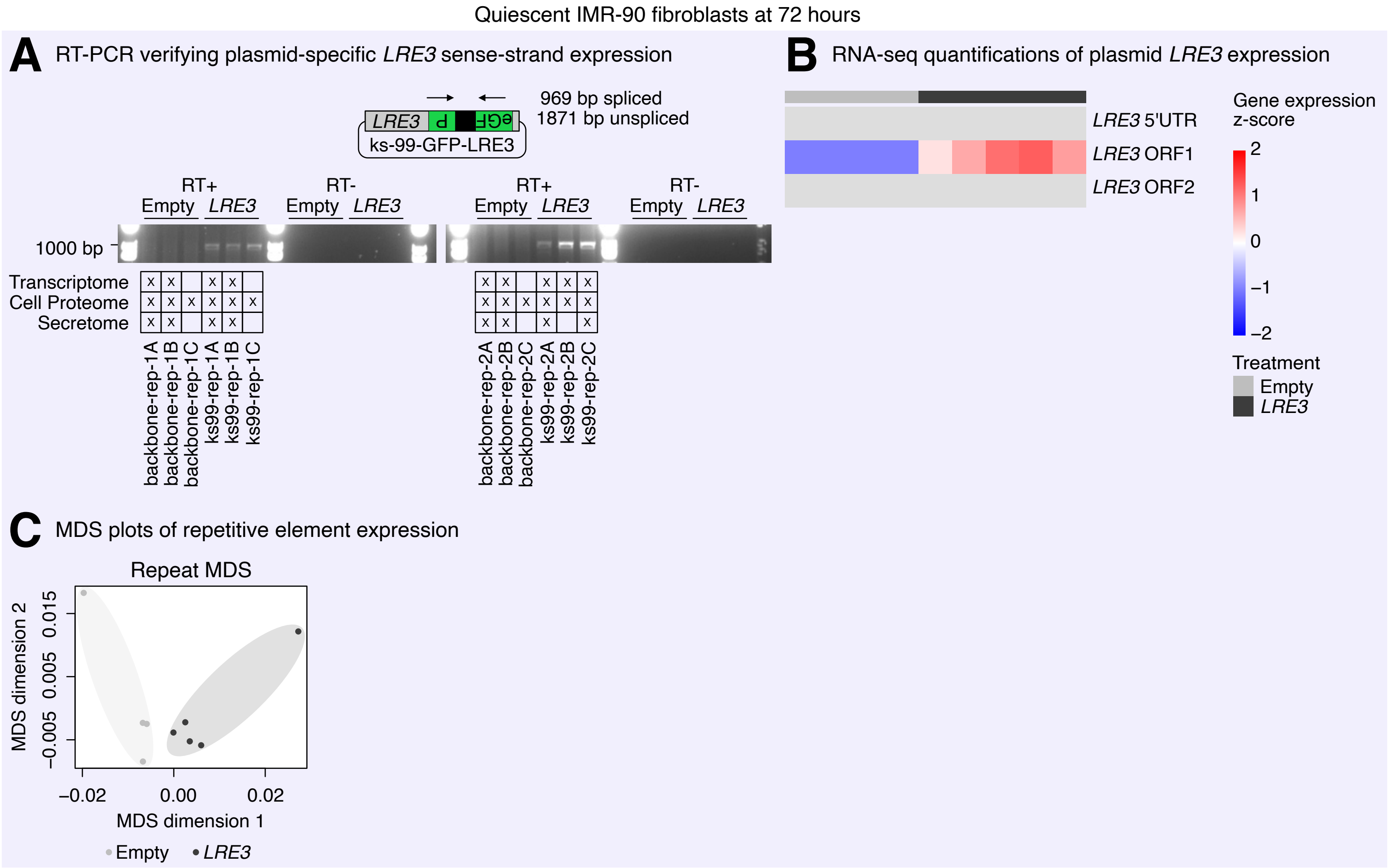

Quiescent IMR-90 fibroblasts at 72 hours
A
RT-PCR verifying plasmid-specific LRE3 sense-strand expression
969 bp spliced
1871 bp unspliced
LRE3
ks-99-GFP-LRE3
RT+
RT-
Empty
LRE3
1000 bp
Transcriptome
Cell Proteome
Secretome
backbone-rep-1A
backbone-rep-1B
backbone-rep-1C
ks99-rep-1A
ks99-rep-1B
ks99-rep-1C
backbone-rep-2A
backbone-rep-2B
backbone-rep-2C
ks99-rep-2A
ks99-rep-2B
ks99-rep-2C
B
RNA-seq quantifications of plasmid LRE3 expression
LRE3 5'UTR
LRE3 ORF1
LRE3 ORF2
Gene expression z-score
2
1
0
−1
−2
Treatment
Empty
LRE3
C
MDS plots of repetitive element expression
Repeat MDS
MDS dimension 2
0.015
0.005
−0.005
−0.02
0.00
0.02
MDS dimension 1
Empty
LRE3

Figure S3

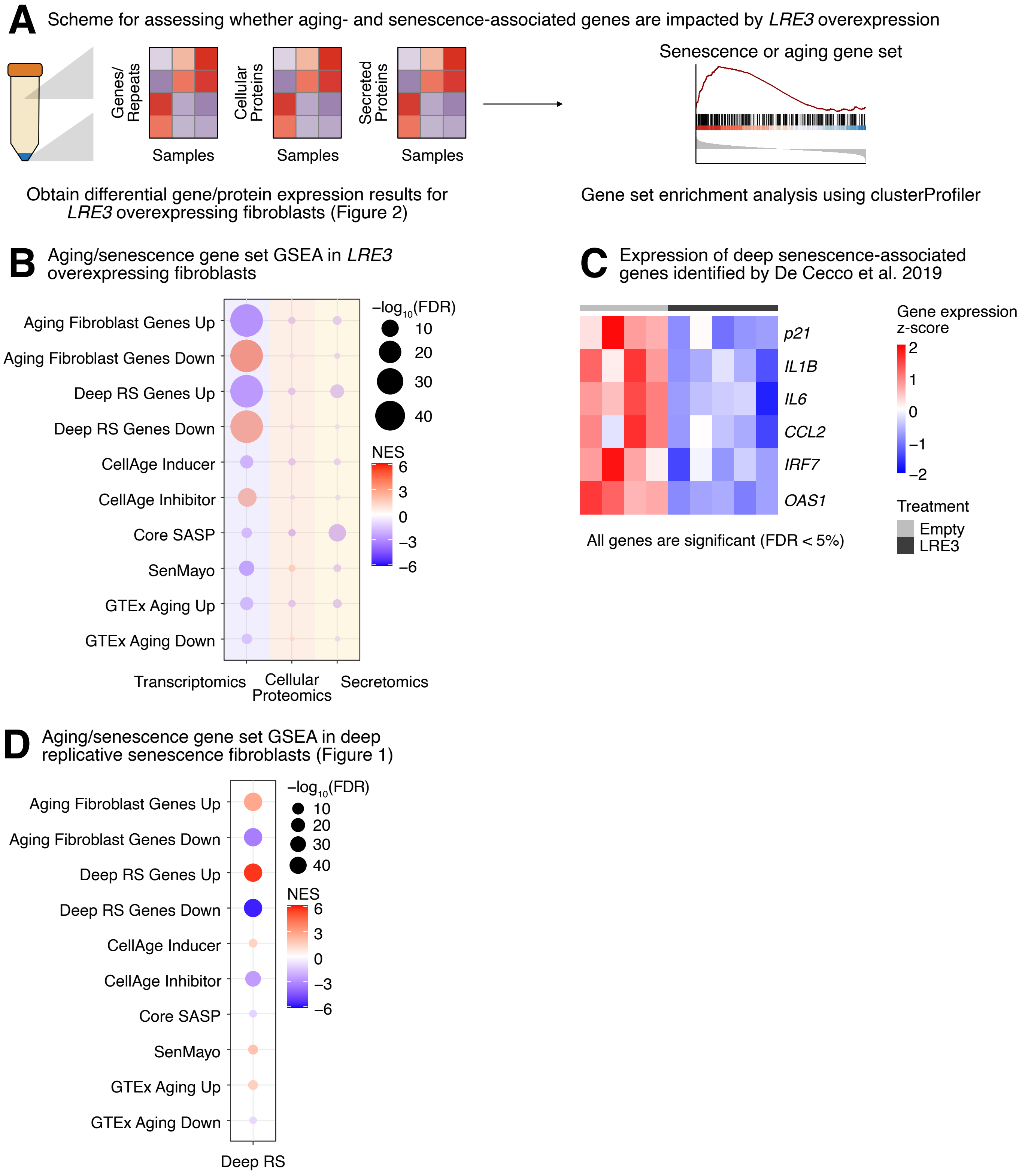
A
Scheme for assessing whether aging- and senescence-associated genes are impacted by LRE3 overexpression
Genes/ Repeats
Cellular Proteins
Secreted Proteins
Samples
Samples
Samples
Senescence or aging gene set
Obtain differential gene/protein expression results for LRE3 overexpressing fibroblasts (Figure 2)
Gene set enrichment analysis using clusterProfiler
B
Aging/senescence gene set GSEA in LRE3 overexpressing fibroblasts
Aging Fibroblast Genes Up
Aging Fibroblast Genes Down
Deep RS Genes Up
Deep RS Genes Down
CellAge Inducer
CellAge Inhibitor
Core SASP
SenMayo
GTEx Aging Up
GTEx Aging Down
Transcriptomics
Cellular Proteomics
Secretomics
−log10(FDR)
10
20
30
40
NES
6
3
0
−3
−6
C
Expression of deep senescence-associated genes identified by De Cecco et al. 2019
p21
IL1B
IL6
CCL2
IRF7
OAS1
All genes are significant (FDR < 5%)
Gene expression z-score
2
1
0
−1
−2
Treatment
Empty
LRE3
D
Aging/senescence gene set GSEA in deep replicative senescence fibroblasts (Figure 1)
Aging Fibroblast Genes Up
Aging Fibroblast Genes Down
Deep RS Genes Up
Deep RS Genes Down
CellAge Inducer
CellAge Inhibitor
Core SASP
SenMayo
GTEx Aging Up
GTEx Aging Down
Deep RS
−log10(FDR)
10
20
30
40
NES
6
3
0
−3
−6

Figure S4

**A** Flow cytometry gating strategy for cell cycle assay (empty vector example)

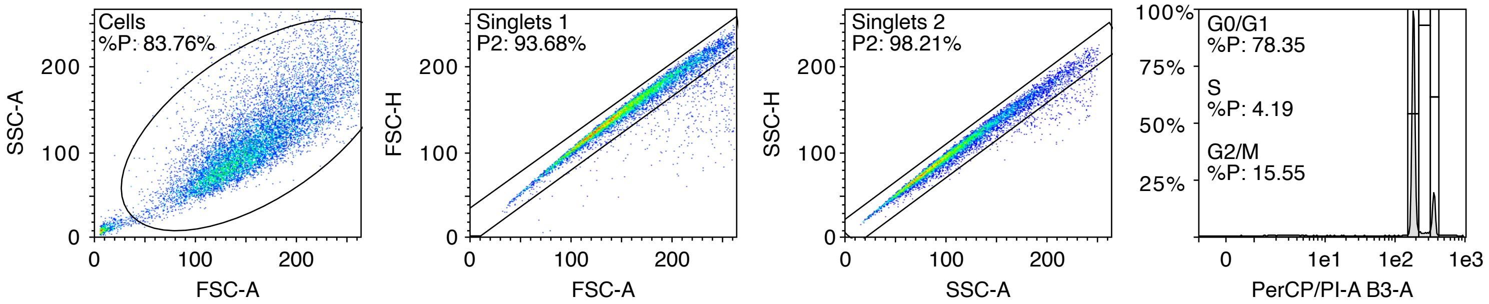

Figure S5

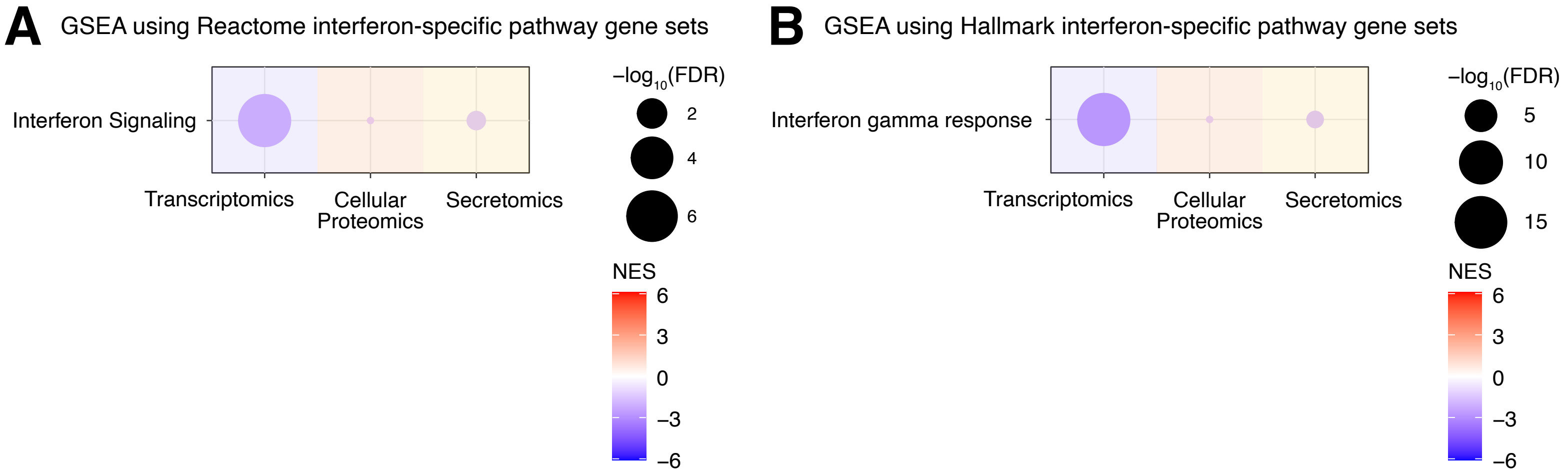
A
GSEA using Reactome interferon-specific pathway gene sets
Interferon Signaling
Transcriptomics
Cellular Proteomics
Secretomics
−log10(FDR)
2
4
6
NES
6
3
0
−3
−6
B
GSEA using Hallmark interferon-specific pathway gene sets
Interferon gamma response
Transcriptomics
Cellular Proteomics
Secretomics
−log10(FDR)
5
10
15
NES
6
3
0
−3
−6

Figure S6

IMR-90 fibroblasts in rich media at 24 hours

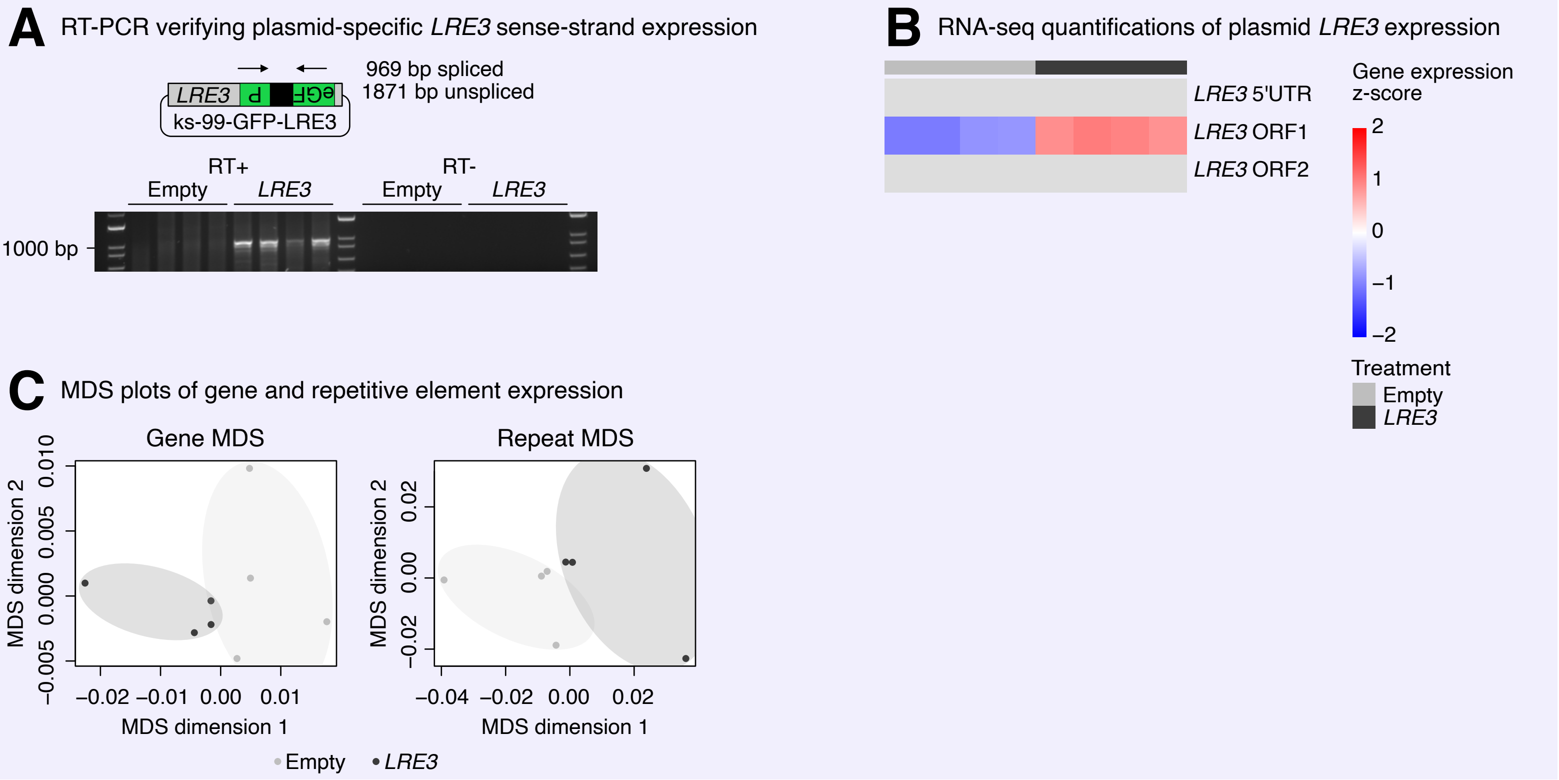

Figure S7

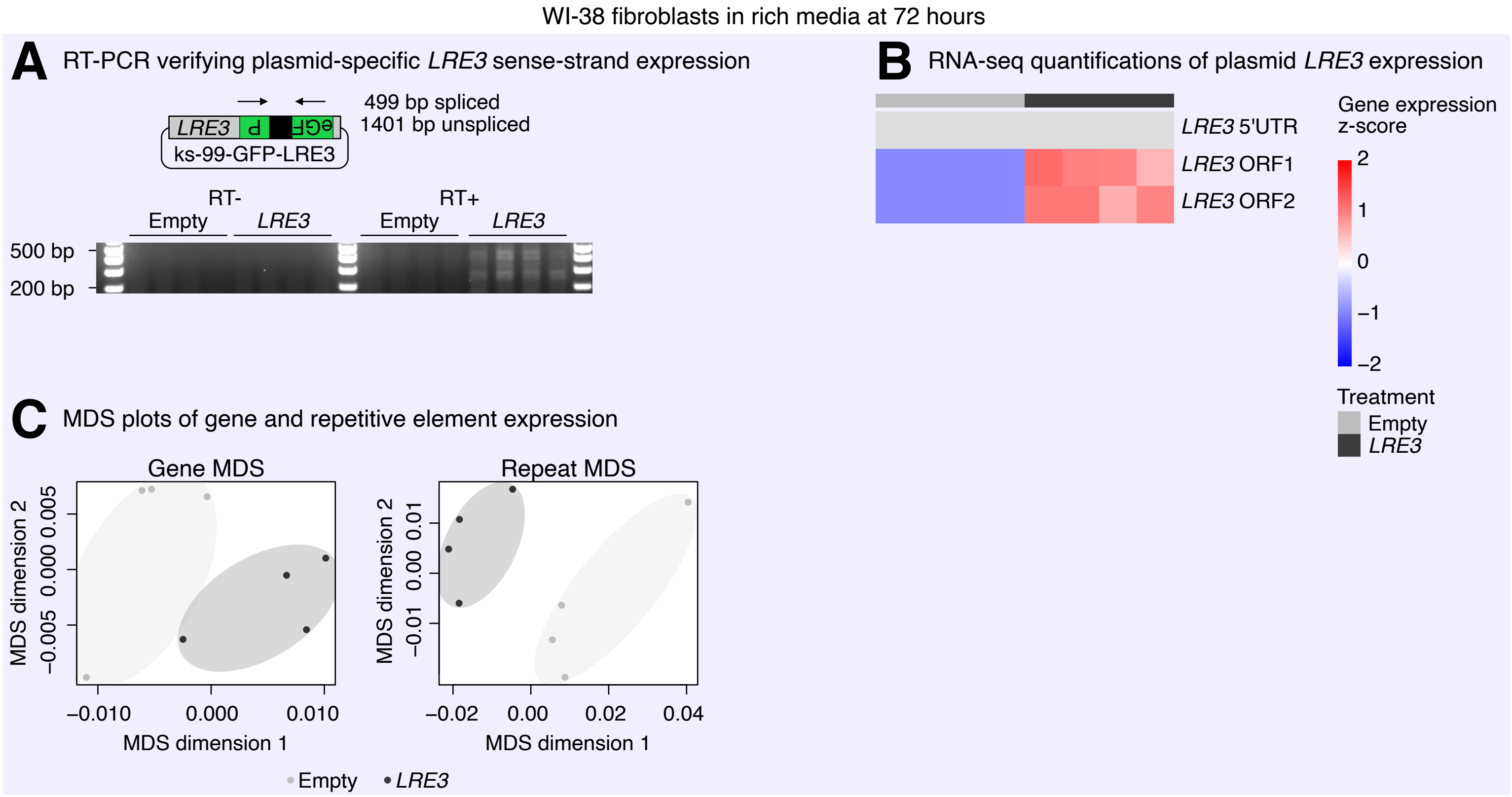
WI-38 fibroblasts in rich media at 72 hours
A RT-PCR verifying plasmid-specific LRE3 sense-strand expression
LRE3
ks-99-GFP-LRE3
499 bp spliced
1401 bp unspliced
RT-
RT+
Empty
LRE3
Empty
LRE3
500 bp
200 bp
B RNA-seq quantifications of plasmid LRE3 expression
LRE3 5'UTR
LRE3 ORF1
LRE3 ORF2
Gene expression z-score
2
1
0
−1
−2
Treatment
Empty
LRE3
C MDS plots of gene and repetitive element expression
Gene MDS
Repeat MDS
MDS dimension 2
−0.005 0.000 0.005
−0.010 0.000 0.010
MDS dimension 1
−0.01 0.00 0.01
−0.02 0.00 0.02 0.04
Empty
LRE3

### Figure S8

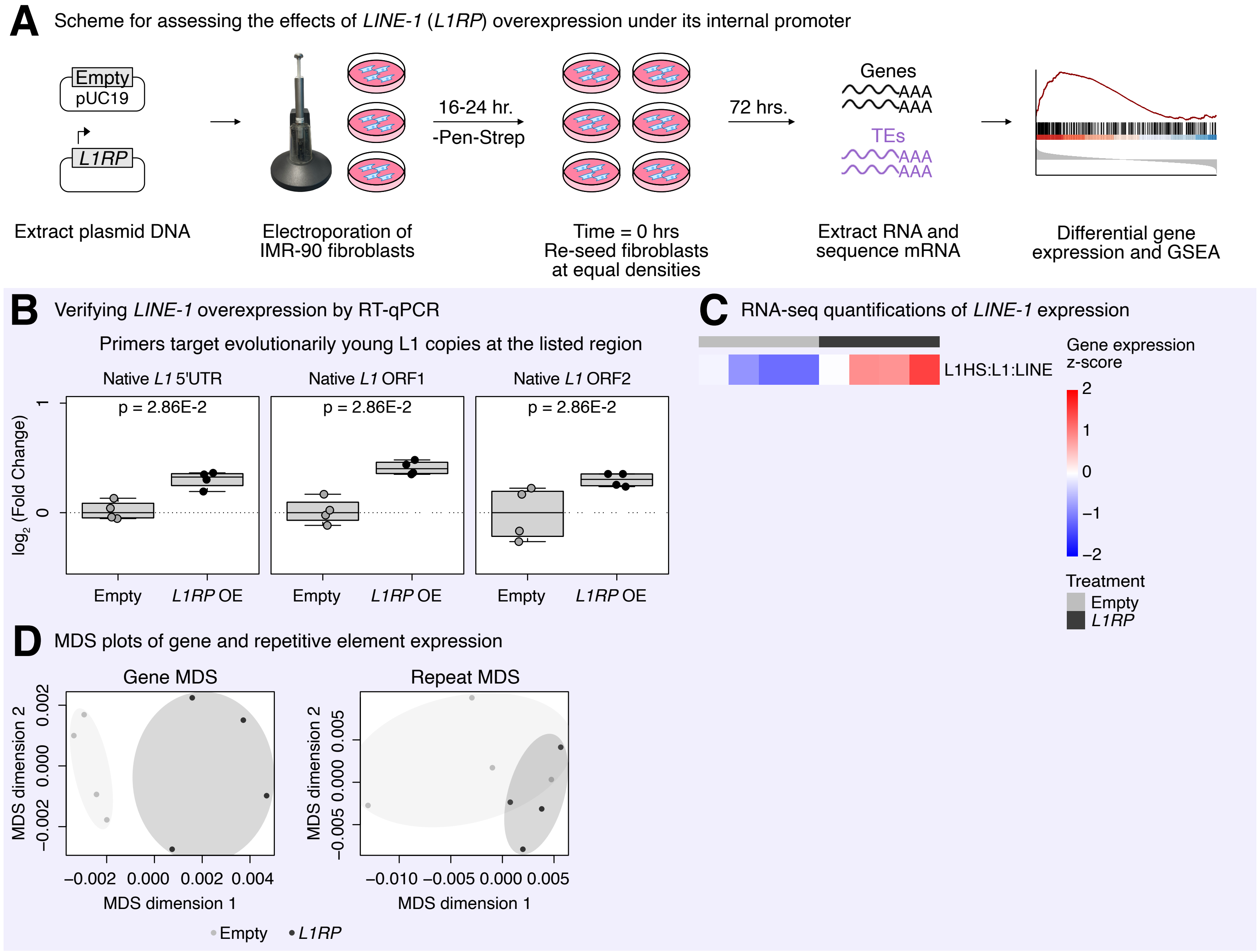
A
Scheme for assessing the effects of LINE-1 (L1RP) overexpression under its internal promoter
Empty
pUC19
L1RP
16-24 hr.
-Pen-Strep
72 hrs.
Genes
AAA
AAA
TEs
AAA
AAA
Extract plasmid DNA
Electroporation of IMR-90 fibroblasts
Time = 0 hrs
Re-seed fibroblasts at equal densities
Extract RNA and sequence mRNA
Differential gene expression and GSEA
B
Verifying LINE-1 overexpression by RT-qPCR
Primers target evolutionarily young L1 copies at the listed region
Native L1 5'UTR
Native L1 ORF1
Native L1 ORF2
p = 2.86E-2
p = 2.86E-2
p = 2.86E-2
log2 (Fold Change)
1
0
Empty
L1RP OE
Empty
L1RP OE
Empty
L1RP OE
C
RNA-seq quantifications of LINE-1 expression
L1HS:L1:LINE
Gene expression z-score
2
1
0
−1
−2
Treatment
Empty
L1RP
D
MDS plots of gene and repetitive element expression
Gene MDS
Repeat MDS
MDS dimension 2
−0.002
0.000
0.002
MDS dimension 2
−0.005
0.000
0.005
−0.002
0.000
0.002
0.004
−0.010
−0.005
0.000
0.005
MDS dimension 1
MDS dimension 1
Empty
L1RP

Figure S9

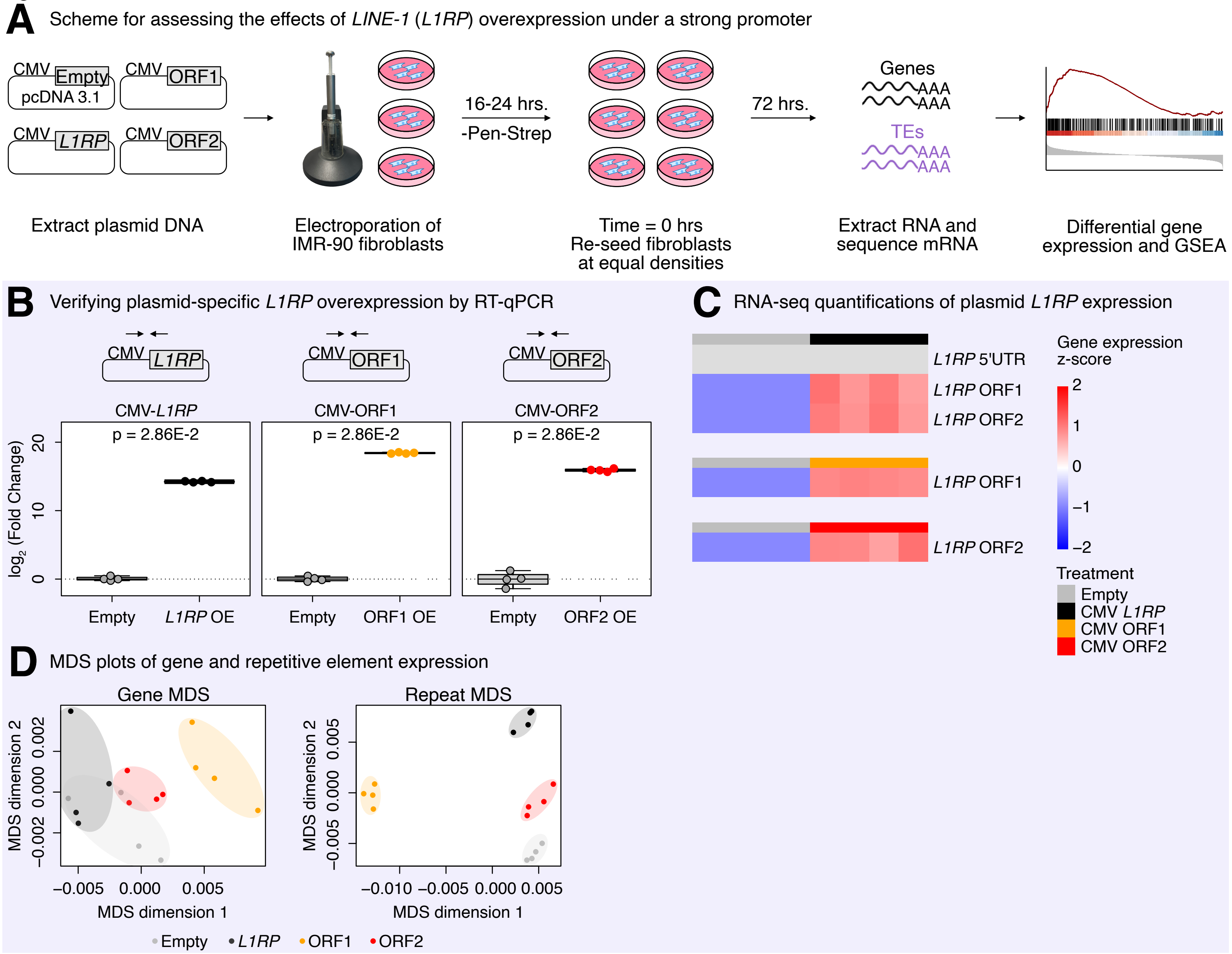
A Scheme for assessing the effects of LINE-1 (L1RP) overexpression under a strong promoter
CMV Empty pcDNA 3.1
CMV ORF1
CMV L1RP
CMV ORF2
16-24 hrs.
-Pen-Strep
72 hrs.
Genes
AAA
AAA
TEs
AAA
AAA
Extract plasmid DNA
Electroporation of IMR-90 fibroblasts
Time = 0 hrs
Re-seed fibroblasts at equal densities
Extract RNA and sequence mRNA
Differential gene expression and GSEA
B Verifying plasmid-specific L1RP overexpression by RT-qPCR
CMV L1RP
CMV ORF1
CMV ORF2
CMV-L1RP
CMV-ORF1
CMV-ORF2
p = 2.86E-2
p = 2.86E-2
p = 2.86E-2
log2 (Fold Change)
0
10
20
Empty
L1RP OE
Empty
ORF1 OE
Empty
ORF2 OE
C RNA-seq quantifications of plasmid L1RP expression
L1RP 5'UTR
L1RP ORF1
L1RP ORF2
L1RP ORF1
L1RP ORF2
Gene expression z-score
2
1
0
−1
−2
Treatment
Empty
CMV L1RP
CMV ORF1
CMV ORF2
D MDS plots of gene and repetitive element expression
Gene MDS
MDS dimension 2
−0.002
0.000
0.002
−0.005
0.000
0.005
MDS dimension 1
Repeat MDS
MDS dimension 2
−0.005
0.000
0.005
−0.010
−0.005
0.000
0.005
MDS dimension 1
Empty
L1RP
ORF1
ORF2

Figure S10

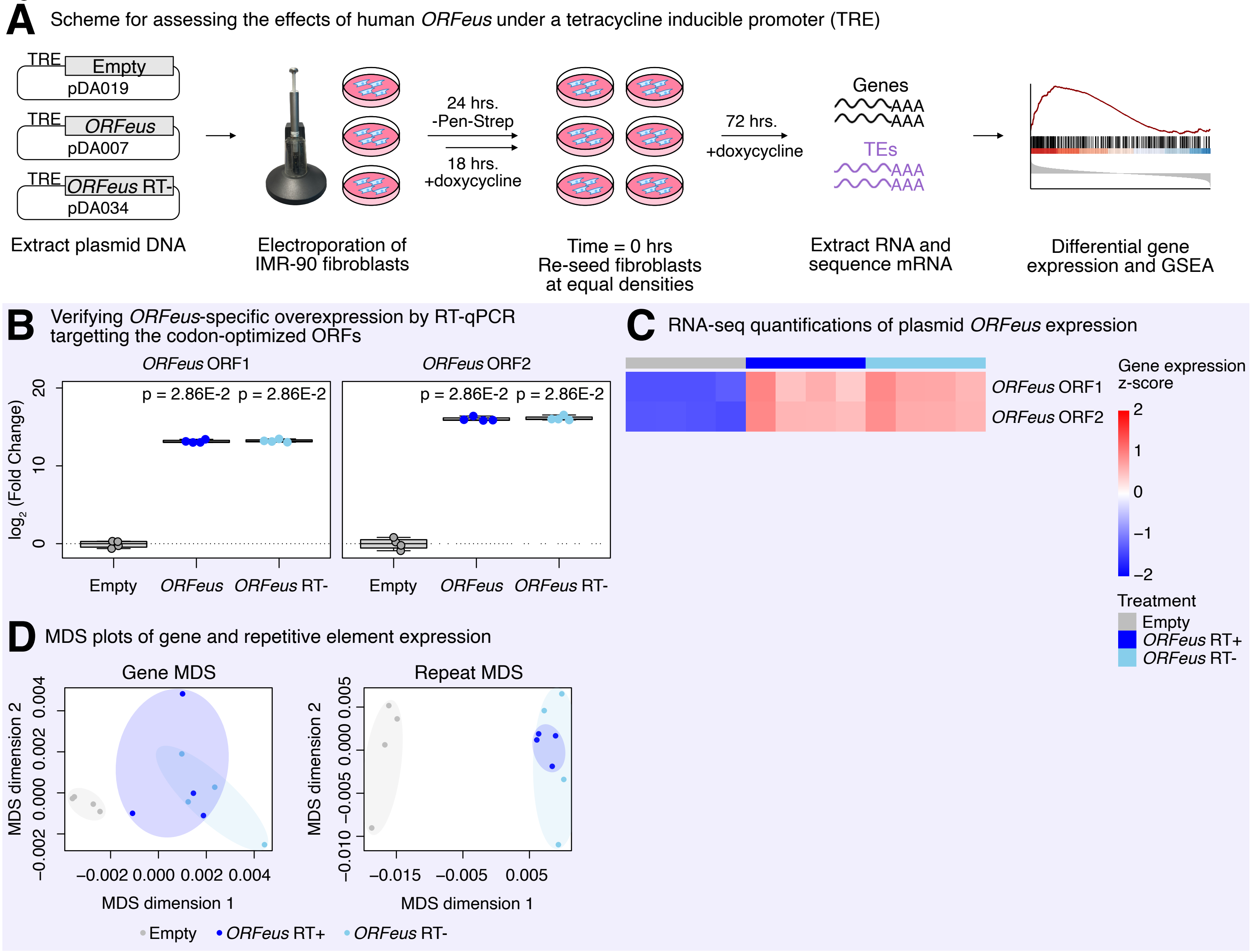
A
Scheme for assessing the effects of human ORFeus under a tetracycline inducible promoter (TRE)
TRE
Empty
pDA019
TRE
ORFeus
pDA007
TRE
ORFeus RT-
pDA034
Extract plasmid DNA
Electroporation of IMR-90 fibroblasts
24 hrs.
-Pen-Strep
18 hrs.
+doxycycline
Time = 0 hrs
Re-seed fibroblasts at equal densities
72 hrs.
+doxycycline
Genes
AAA
AAA
TEs
AAA
AAA
Extract RNA and sequence mRNA
Differential gene expression and GSEA
B
Verifying ORFeus-specific overexpression by RT-qPCR targetting the codon-optimized ORFs
ORFeus ORF1
ORFeus ORF2
p = 2.86E-2
p = 2.86E-2
p = 2.86E-2
p = 2.86E-2
log2 (Fold Change)
20
10
0
Empty
ORFeus
ORFeus RT-
Empty
ORFeus
ORFeus RT-
C
RNA-seq quantifications of plasmid ORFeus expression
ORFeus ORF1
ORFeus ORF2
Gene expression z-score
2
1
0
−1
−2
Treatment
Empty
ORFeus RT+
ORFeus RT-
D
MDS plots of gene and repetitive element expression
Gene MDS
MDS dimension 2
−0.002 0.000 0.002 0.004
MDS dimension 1
−0.002 0.000 0.002 0.004
Repeat MDS
MDS dimension 2
−0.010 −0.005 0.000 0.005
MDS dimension 1
−0.015 −0.005 0.005
Empty
ORFeus RT+
ORFeus RT-

### Figure S11

HeLa cells in rich media at 48 hours

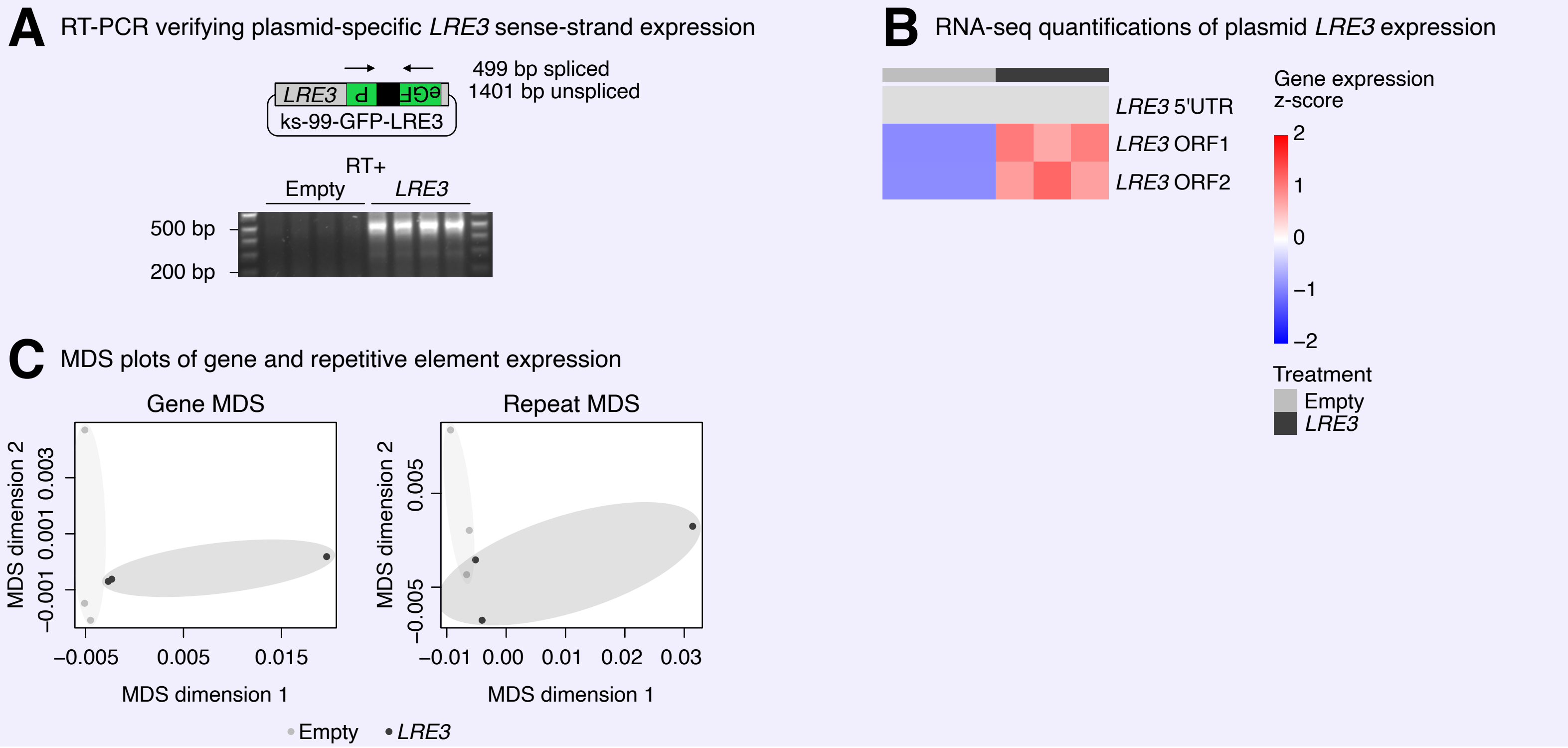

Figure S12

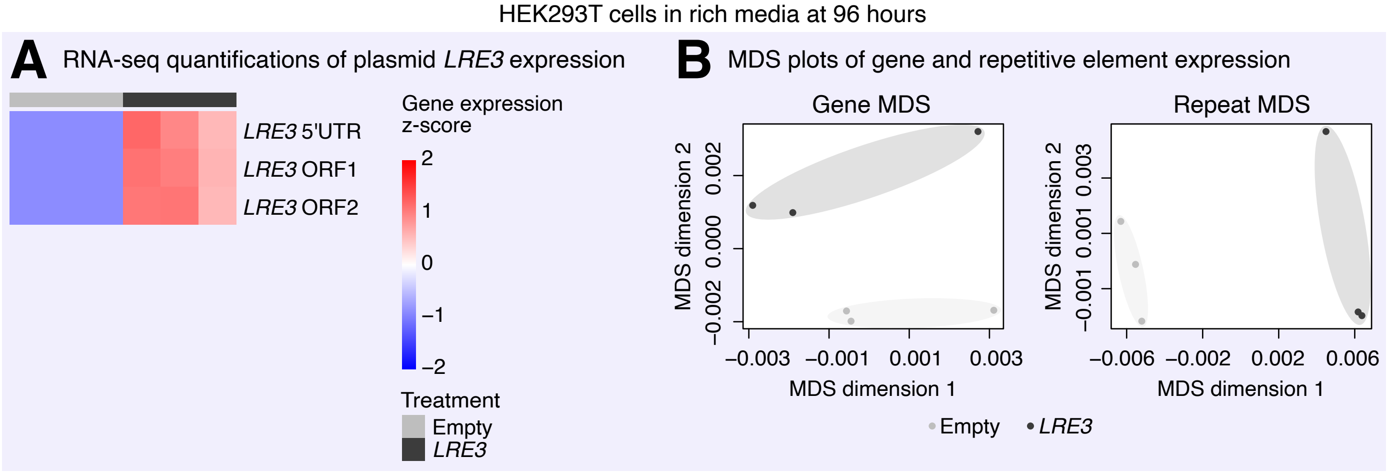
HEK293T cells in rich media at 96 hours
A
RNA-seq quantifications of plasmid LRE3 expression
LRE3 5'UTR
LRE3 ORF1
LRE3 ORF2
Gene expression z-score
2
1
0
−1
−2
Treatment
Empty
LRE3
B
MDS plots of gene and repetitive element expression
Gene MDS
MDS dimension 2
−0.002 0.000 0.002
−0.003 −0.001 0.001 0.003
MDS dimension 1
Repeat MDS
MDS dimension 2
−0.001 0.001 0.003
−0.006 −0.002 0.002 0.006
MDS dimension 1
Empty
LRE3

Figure S13

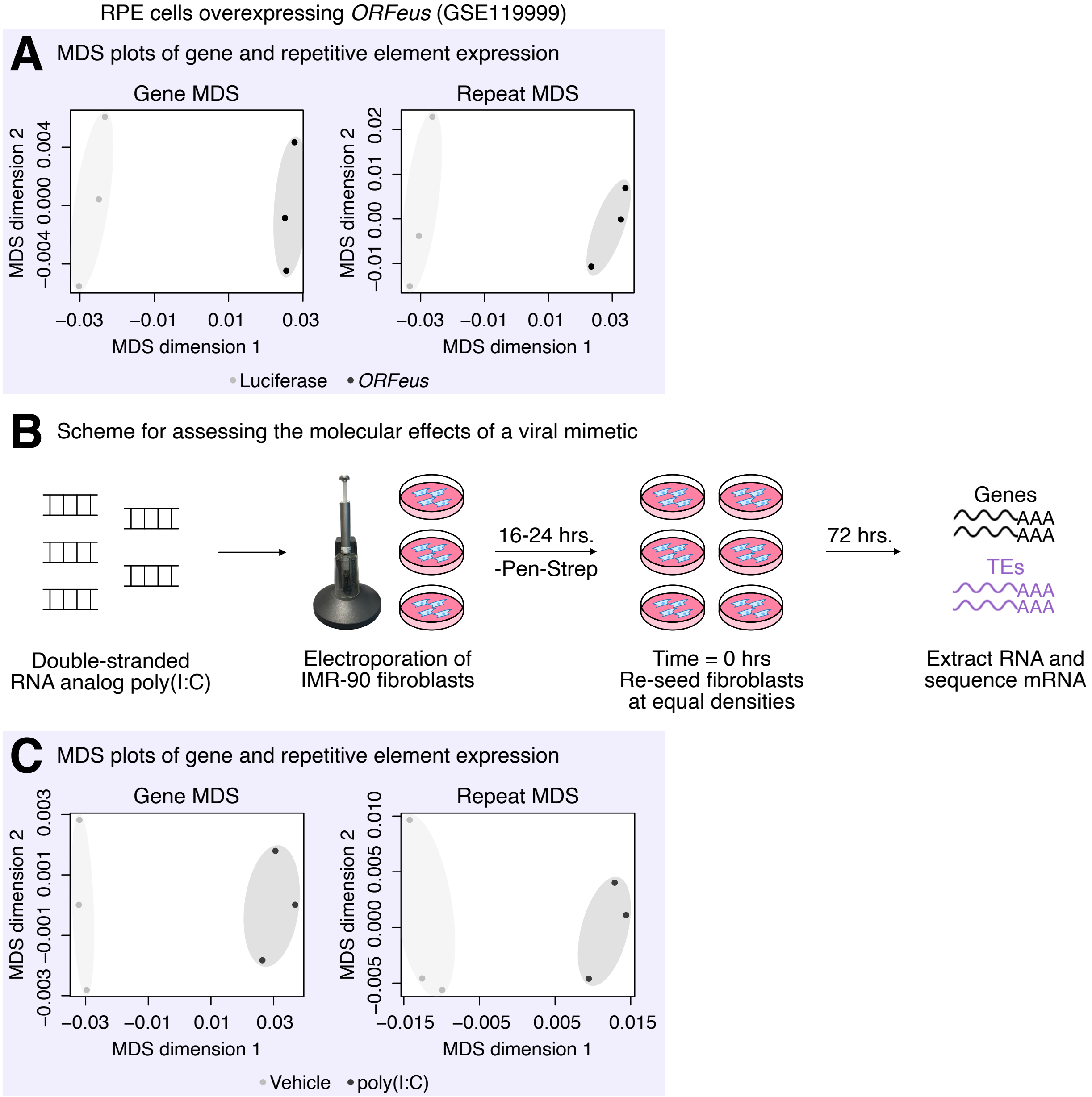
RPE cells overexpressing ORFeus (GSE119999)
A
MDS plots of gene and repetitive element expression
Gene MDS
Repeat MDS
MDS dimension 2
MDS dimension 1
−0.004 0.000 0.004
−0.01 0.00 0.01 0.02
−0.03 −0.01 0.01 0.03
Luciferase
ORFeus
B
Scheme for assessing the molecular effects of a viral mimetic
Double-stranded RNA analog poly(I:C)
Electroporation of IMR-90 fibroblasts
16-24 hrs.
-Pen-Strep
Time = 0 hrs
Re-seed fibroblasts at equal densities
72 hrs.
Genes
AAA
AAA
TEs
AAA
AAA
Extract RNA and sequence mRNA
C
MDS plots of gene and repetitive element expression
Gene MDS
Repeat MDS
MDS dimension 2
MDS dimension 1
−0.003 −0.001 0.001 0.003
−0.005 0.000 0.005 0.010
−0.03 −0.01 0.01 0.03
−0.015 −0.005 0.005 0.015
Vehicle
poly(I:C)

Figure S14

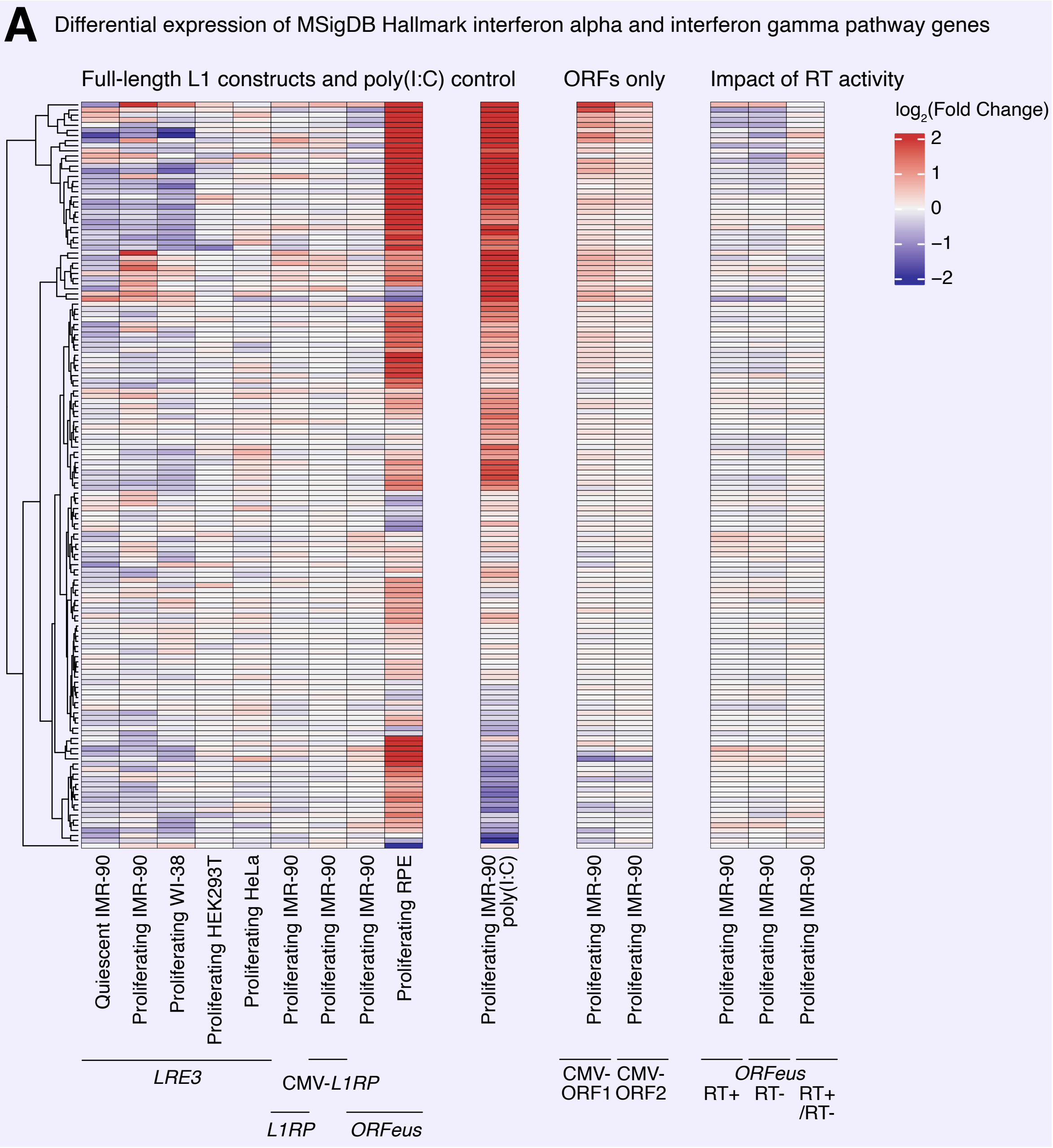
A
Differential expression of MSigDB Hallmark interferon alpha and interferon gamma pathway genes
Full-length L1 constructs and poly(I:C) control
ORFs only
Impact of RT activity
log2(Fold Change)
2
1
0
−1
−2
Quiescent IMR-90
Proliferating IMR-90
Proliferating WI-38
Proliferating HEK293T
Proliferating HeLa
Proliferating IMR-90
Proliferating IMR-90
Proliferating IMR-90
Proliferating RPE
Proliferating IMR-90 poly(I:C)
Proliferating IMR-90
Proliferating IMR-90
Proliferating IMR-90
Proliferating IMR-90
Proliferating IMR-90
LRE3
CMV-L1RP
L1RP
ORFeus
CMV-ORF1
CMV-ORF2
ORFeus
RT+
RT-
RT+/RT-
